# CryoEM structures of human CMG - ATPγS - DNA and CMG - AND-1 complexes

**DOI:** 10.1101/2020.01.22.914192

**Authors:** Neil J Rzechorzek, Steven W Hardwick, Vincentius A Jatikusumo, Dimitri Y Chirgadze, Luca Pellegrini

**Affiliations:** Department of Biochemistry, Tennis Court Road, Cambridge CB2 1GA, UK

## Abstract

DNA unwinding in eukaryotic replication is performed by the Cdc45-MCM-GINS (CMG) helicase. Although the CMG architecture has been elucidated, its mechanism of DNA unwinding and replisome interactions remain poorly understood. Here we report the cryoEM structure at 3.3 Å of human CMG bound to fork DNA and the ATP-analogue ATPγS. Eleven nucleotides of single-stranded (ss) DNA are bound within the C-tier of MCM2-7 AAA+ ATPase domains. All MCM subunits contact DNA, from MCM2 at the 5′-end to MCM5 at the 3′-end of the DNA spiral, but only MCM6, 4, 7 and 3 make a full set of interactions. DNA binding correlates with nucleotide occupancy: five MCM subunits are bound to either ATPγS or ADP, whereas the apo MCM2-5 interface remains open. We further report the cryoEM structure of human CMG bound to the replisome hub AND-1 (CMGA). The AND-1 trimer uses one β-propeller domain of its trimerisation region to dock onto the side of the helicase assembly formed by Cdc45 and GINS. In the resulting CMGA architecture, the AND-1 trimer is closely positioned to the fork DNA while its CIP (Ctf4-interacting peptide)-binding helical domains remain available to recruit partner proteins.

---

Accurate and faithful duplication of our chromosomal DNA in preparation for mitosis is essential for cellular life (Michael O’Donnell, Langston, & Stillman, 2013). DNA synthesis in S-phase is a highly complex biochemical process carried out by the replisome, a large and dynamic multi-protein assembly of about thirty core components (Bell & Labib, 2016). The replisome contains all necessary enzymatic activities for copying the genetic information encoded in the parental DNA, as well as non-enzymatic factors that guarantee efficient DNA synthesis under normal conditions and during replicative stress.

Central to the replication process is the physical separation of the parental strands of DNA, to allow the templated polymerisation of new leading and lagging strands, according to the semi-discontinuous model of DNA replication. Replicative DNA helicases form hexameric rings that thread single-stranded DNA through their ring channel and achieve unwinding of double-stranded (ds) DNA by a process of strand exclusion (Lyubimov, Strycharska, & Berger, 2011; Michael E O’Donnell & Li, 2018). Each helicase subunit consists of an N-terminal domain and a C-terminal ATPase domain, which form a double stack of N-tier and C-tier rings. ATP binding takes place at the subunit interface and its hydrolysis requires residues from both subunits. Processive strand separation results from the allosteric coupling of ATP hydrolysis to concerted movements of the DNA-binding elements that line the ring pore in each subunit, as first shown for the replicative viral E1 DNA helicase (Enemark & Joshua-Tor, 2006) and the hexameric Rho RNA helicase (Thomsen & Berger, 2009).

Unwinding of parental DNA in eukaryotic cells is performed by the 11-subunit Cdc45-MCM-GINS assembly or CMG (Moyer, Lewis, & Botchan, 2006). The six MCM proteins, MCM2-7, belong to the AAA+ family of ATPases and form a hetero-hexameric ring that translocates on single-strand DNA (Abid Ali & Costa, 2016). In the absence of DNA substrate, MCM2-7 adopts predominantly an open spiral conformation; the co-factors Cdc45 and GINS — a hetero-tetramer of Psf1-3 and Sld5 — bind to the MCM5-2 interface and lock MCM2-7 into the closed-ring conformation required for robust DNA unwinding (Costa et al., 2011). In vivo, a multi-step process operates to assemble and activate the eukaryotic CMG DNA helicase at the start of S-phase. In the model system budding yeast, CMG activation involves loading of the MCM2-7 proteins at origin DNA as an inactive double hexamer, which is then activated by phosphorylation-dependent recruitment of the Cdc45 and GINS cofactors, intervention of MCM10, and ATP hydrolysis (Douglas, Ali, Costa, & Diffley, 2018). Activation yields two CMG assemblies that segregate on opposite template DNA strands and move past each other to establish two independent replication forks (Georgescu et al., 2017). The translocating CMG tracks along the leading-strand template in the 3′-to-5′ direction (Fu et al., 2011), with the N-tier ring of MCM2-7 at the leading edge of the advancing helicase (Georgescu et al., 2017). Strand separation is proposed to be achieved by a modified version of steric exclusion, whereby the lagging strand penetrates the N-tier of the CMG before separation (Langston & O’Donnell, 2017).

The mechanism of translocation by which the CMG couples ATP hydrolysis to processive DNA unwinding is the current focus of intense research efforts. Based on structural analysis of bacteriophage, viral and bacterial systems (Enemark & Joshua-Tor, 2006; Gao et al., 2019; Itsathitphaisarn, Wing, Eliason, Wang, & Steitz, 2012; Singleton, Sawaya, Ellenberger, & Wigley, 2000) a consensus has emerged for a sequential rotary mechanism of DNA unwinding by replicative DNA helicases. In this mechanism, ATP is sequentially hydrolysed by successive ring subunits so that each ring position cycles through ATP, ADP and apo states. In turn, the ATP state determines allosterically the position of the DNA-binding loops, that adopt a staircase arrangement matching the DNA spiral bound within the ring pore. The sequential hydrolysis of ATP around the ring causes the coordinated motion of the DNA-binding loops, resulting in translocation of the DNA substrate through the ring.

A complicating feature when trying to analyse CMG translocation is that, unlike the homo-hexameric helicases of simpler organisms, the MCM2-7 motor of the CMG is a hetero-hexamer of six related but distinct subunits (Bochman, Bell, & Schwacha, 2008). Indeed, biochemical measures of DNA unwinding by purified fly CMG showed that ATP binding and hydrolysis are not equally important at all MCM ring interfaces (Eickhoff et al., 2019; Ilves, Petojevic, Pesavento, & Botchan, 2010). Furthermore, biological evidence in yeast shows that the importance of DNA binding is different among MCM subunits (Lam et al., 2013; Ramey & Sclafani, 2014). Recent cryoEM analyses of yeast CMG have led to the proposal of alternative translocation mechanisms, based on ‘pumpjack’ or ‘inchworm’ movements of the N- and C-tier of the MCM ring (Abid Ali et al., 2016; Yuan et al., 2016). A recent structural study of the fly CMG in conditions of DNA-fork unwinding (Eickhoff et al., 2019) imaged four distinct states of the helicase; the states formed the basis for an asymmetric model of DNA unwinding that accounted for the different roles of the MCM2-7 subunits in translocation.

The critical insights provided by these initial landmark studies have not been sufficient to settle the important issue of the mechanism of DNA translocation by the CMG, and therefore further structural investigations are needed. It is especially important to obtain high-resolution cryoEM maps that will allow the determination of accurate atomic models of the helicase bound to fork DNA substrates, to elucidate unambiguously key aspects of the mechanism of translocation on DNA such as the protein-DNA interface and the geometry of the ATP-binding sites. Equally important is to obtain high-resolution information on the interactions of the CMG with other core replisome components. Furthermore, published structural analyses focused on CMGs from simpler model systems such as yeast or Drosophila, and no structural evidence is currently available for vertebrate CMG.

Here we report the cryoEM structure at 3.3 Å of human CMG bound to a fork DNA substrate in the presence of ATPγS. We also present the cryoEM structure of human CMG bound to AND-1, a core replisome component that acts as a platform for recruitment of replisome components to the replication fork. Unique features captured in our structures provide insights into DNA translocation and formation of larger replisome assemblies by the human CMG helicase.

## RESULTS

### Expression and purification of human CMG-ATPγS-DNA

To maximise our chances of producing correctly-assembled human CMG, we used transient transfection of suspension-free HEK293 cells with a plasmid system encoding all 11 subunits of the CMG assembly. After co-expression of MCM2-7, Cdc45 and GINS, human CMG was purified by Ni^2+^- and Streptactin-affinity chromatography (**Supplementary figure 1A**). A large endogenous protein that co-purified at sub-stoichiometric levels with the CMG over the two-step purification was identified as AND-1, a known replisome component. AND-1 co-purification indicates a tight constitutive association with the CMG in the human replisome, in agreement with the known association of AND-1’s orthologue Ctf4 with the yeast CMG (Gambus et al., 2006).

To capture a high-resolution snapshot of human CMG poised to translocate on a fork DNA substrate, we decided to use the ATP analogue ATPγS. Streptactin-bound CMG was incubated with buffer containing a fork DNA substrate and ATPγS before elution with desthiobiotin (**Supplementary figure 1B**). The DNA consisted of a 40 bp duplex region with 30 nt tails and resembled closely a fork DNA that had been designed to measure CMG’s helicase activity, with a 3*′* polydT tail for helicase loading in the correct orientation for fork unwinding and a 5*′* GC-rich tail that inhibits helicase binding (Petojevic et al., 2015).

CryoEM data were collected on a Titan Krios operating at 300 keV using a K2 Summit detector and processed with Relion-3 (Scheres, 2012). After 2D and 3D classification and refinement, we obtained a 3.29 Å map of CMG-DNA-ATPγS from a set of 213,527 particles, and a 3.41 Å map of the CMG C-tier, comprising the ring of AAA+ ATPase domains, after masking of the N-tier (**Supplementary figures 2 and 3**). Both maps were used to build a molecular model of CMG--ATPγS-DNA. The excellent quality and high resolution of the map allowed an accurate description at atomic level of the protein-DNA interface and ATP-binding sites of the human CMG (**Figure 1A**).

**Figure 1.**
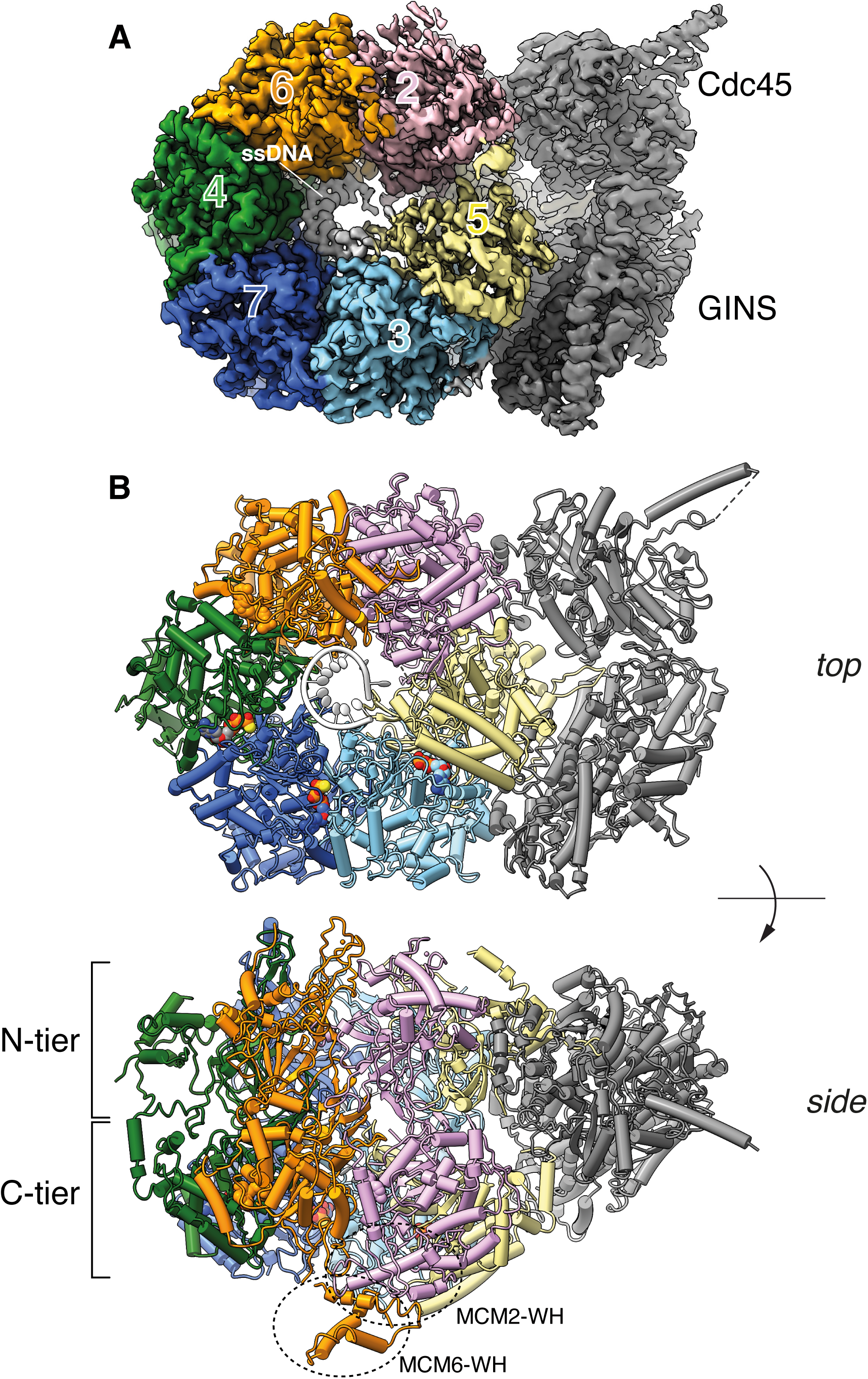
Structure of the human CMG-ATPγS-DNA DNA helicase. **A** 3.3Å cryoEM map of CMG-ATPγS-DNA. MCM2-7 are coloured according to chain: MCM2 is pink, MCM3 cyan, MCM4 green, MCM5 yellow, MCM6 orange and MCM7 blue, following the colour scheme of Eickhoff and colleagues (Eickhoff et al., 2019). The DNA is coloured white, and the Cdc45 and GINS grey. **B** Top and side views of a ribbon model of the atomic structure of CMG-ATPγS-DNA. The N- and C-tier of the CMG are highlighted in the side view, as well as the location of MCM2 and MCM6’s C-terminal WH domains. Colouring as in panel A.

### Overall structure

The 11-subunit assembly of the human CMG shows the familiar architecture first demonstrated for the yeast and drosophila CMG (Costa et al., 2011; Georgescu et al., 2017): a two-tiered ring of MCM2-7 proteins, with the Cdc45 and GINS coactivators bound together to the N-tier portion of MCM2, MCM5 and MCM3, so that Cdc45 faces the MCM2-5 interface of ATPase domains (**Figure 1B**). Each AAA+ ATPase domain contains a nucleotide-binding site at the subunit interface in the C-tier ring, and interacts with DNA via two β-hairpin loops named pre-sensor-1 (PS1) and helix-2 insert (H2I) (Iyer, Leipe, Koonin, & Aravind, 2004) that line the pore of the C-tier ring (**Supplementary figure 4A**). Flexible anchorage between MCM subunits is provided by a domain-swapped helix in each ATPase domain, which tethers each MCM to its neighbour subunit (**Supplementary figure 4B**).

A continuous chain of eleven thymidine nucleotides is bound in a right-handed B-form spiral within the C-tier channel of ATPase domains. The single-stranded (ss) DNA contacts all 6 MCM subunits, from MCM2 with its 5′-end to MCM5 with the 3′-end, and thus traverses all MCM interfaces except MCM2-5 (**Figure 2A, B**). No clear density is visible within the N-tier of the CMG for either single-stranded DNA or the double-strand portion of our fork DNA substrate. The likely explanation for this observation is that the slowly-hydrolysable ATPγS nucleotide has permitted the engagement of the CMG helicase with the leading-strand portion of the fork DNA substrate but prevented its translocation to the ss-dsDNA nexus.

**Figure 2.**
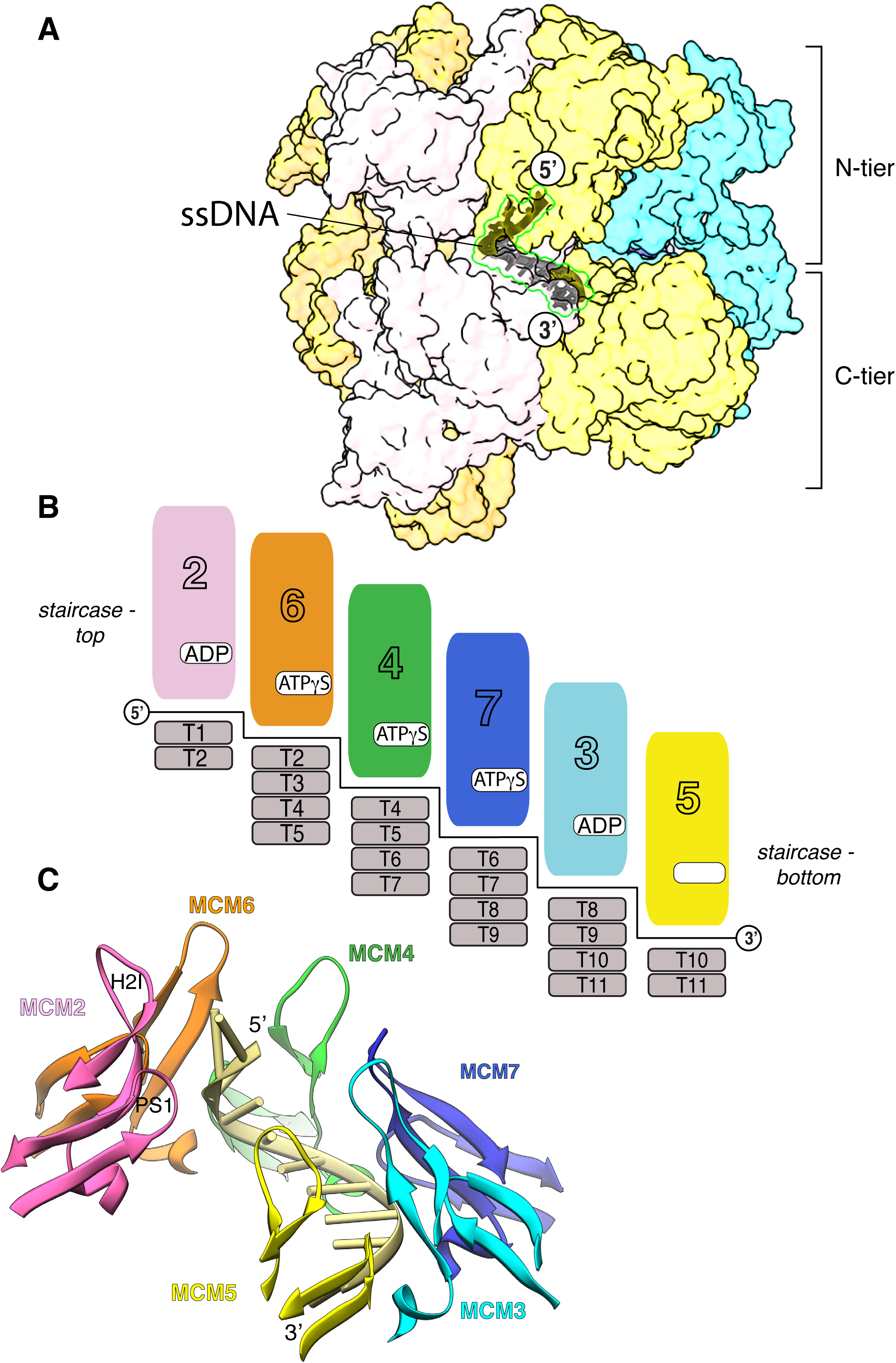
Protein-DNA interactions in the human CMG-ATPγS-DNA complex. MCM chains are coloured coded as in Figure 1. **A** Side view of the CMG-ATPγS-DNA complex, highlighting the position of ssDNA within the MCM C-tier. The MCM subunits are shown as transparent molecular surfaces and the DNA is drawn as a dark grey ribbon. Cdc45 and GINS have been omitted for clarity. **B** Schematic drawing illustrating the DNA footprint and ATP status of each MCM subunit. **C** The staircase configuration of the H2I and PS1 DNA-binding loops around the ssDNA spiral, from MCM2 (pink) at the top (5′-end of the DNA) to MCM5 (yellow) at the bottom (3′-end).

Three of the six MCM interfaces in the ring: MCM6-4, MCM4-7 and MCM7-3, are bound to ATPγS, whereas the MCM3-5 and MCM2-6 interfaces contain ADP as product of ATPγS hydrolysis, while the MCM2-5 interface is empty (**Figure 2B**). In accordance with the apo status of MCM5, the MCM2-5 gate remains ajar and the MCM2-7 C-tier ring adopts a shallow right-handed spiral conformation (**Supplementary Figure 5**).

In addition to their N- and C-tier domains, each MCM subunit contains a smaller C-terminal winged helix (WH) domain. The WH domains of MCM2 and MCM6 were well resolved in the focused C-tier map and could be therefore be modelled in the density (**Figure 1B**). Density for the WH domain of MCM5 could be identified in the lumen of the C-tier pore, but was not of sufficient quality to allow modelling. The similar MCM2 and 6 WH domains sit on the rim of the C-tier and interact with each other with approximate two-fold symmetry. The first 14 proline-rich amino acids of the MCM3 isoform used in our study bind at the interface between the MCM3 N-tier and the GINS Psf3, likely extending and stabilising the GINS-MCM ring interface (**Supplementary Figure 6**).

### DNA binding

In the structure, the ssDNA is embedded within the pore of the C-tier ring (**Figure 2A**). Nine of the eleven nucleotides from the 3′-end of the ssDNA adopt a right-handed spiral conformation that follows closely that of B-form DNA. Four MCM subunits, MCM6, 4, 7 and 3 make an identical set of contacts with DNA, involving both PS1 and H2I loops. MCM6, 4, 7 and 3 interact with four nucleotides each, with a two-nucleotide offset between contiguous subunits (**Figure 2B**). The DNA-binding loops in are arranged in a staircase matching the DNA spiral, from MCM2 at the top of the staircase (5′-end of the DNA) to MCM5 at the bottom (3′-end) (**Figure 2C**). DNA binding correlates with nucleotide occupancy, as nucleotide-bound MCM6, 4, 7 and 3 make a full set of interactions with DNA.

Within each four-nucleotide footprint, an invariant serine at the start of loop H2I (MCM6 S425, MCM4 S539, MCM7 S410, MCM3 S419) is hydrogen bonded to the 5′-terminal phosphate, whilst an invariant lysine in loop PS1 (MCM6 K486, MCM4 K600, MCM7 K471, MCM3 K480) is ion paired to the subsequent phosphate (**Figure 3** and **Supplementary figures 7, 8A**). The role of the invariant PS1 lysine is remarkably similar to that of K506 of the E1 papillomavirus replicative DNA helicase (Enemark & Joshua-Tor, 2006). The ssDNA is kept in close contact with each MCM subunit by two hydrogen bonds between phosphates of the second and third nucleotide in each binding site and main-chain nitrogens of the PS1 residue after the invariant lysine and of a first-strand residue in the H2I hairpin (**Figure 3**).

**Figure 3.**
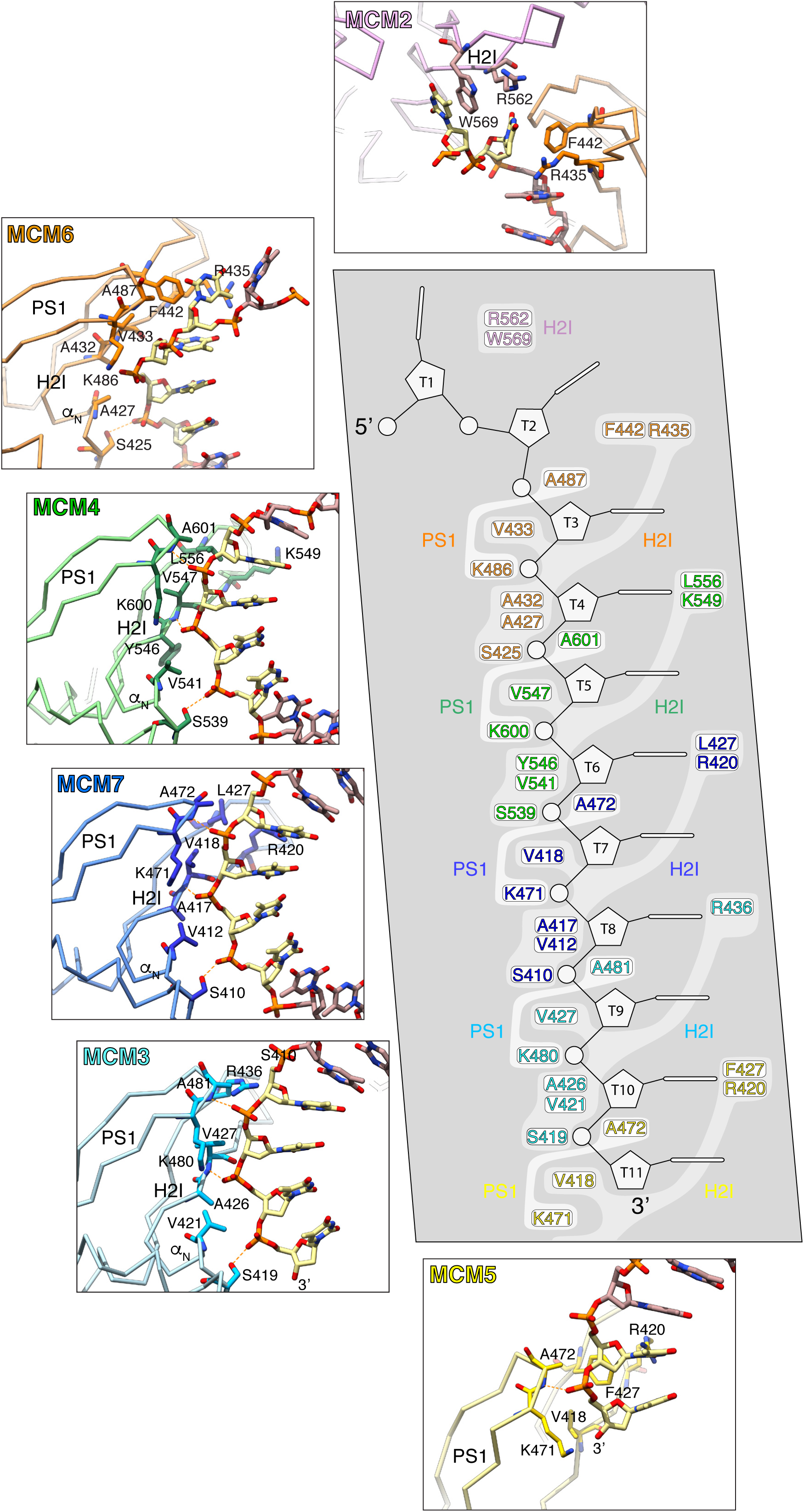
Atomic details of the CMG-DNA interface. The central panel shows a schematic drawing of the 11-nucleotide ssDNA and the MCM residues that interact with DNA in the H2I and PS1 loops of each subunit. In the six surrounding boxes, close-up views of the protein-DNA interface for each MCM subunits are shown. MCM chains are colour-coded as in Figure 1.

Besides these polar contacts, the protein-DNA interface has substantial hydrophobic character: small aliphatic side chains of valine and alanine in both H2I and PS1 loops pack against the ribose-phosphate backbone of the DNA, creating a continuous hydrophobic surface in the C-tier pore that matches the spiral of the DNA (**Figure 3**). In addition to making extensive contacts with the DNA backbone, the MCM subunits use the H2I loop to interact with the bases: a pair of conserved H2I residues, consisting of a basic and an aromatic/hydrophobic amino acid six residues apart ([+]x_6_[Ω/Ψ] motif; +, basic; Ω, aromatic; Ψ, large aliphatic) contact the third and fourth thymidine in each binding site (**Figure 3** and **Supplementary figure 7**).

The B-form spiral of the DNA is interrupted at the MCM6-DNA interface. MCM6 F442 — the aromatic residue in the [+]x_6_[Ω/Ψ] motif — unstacks the second and third nucleotide from the 5′-end of the DNA by inserting its side chain between the thymine bases (**Figure 3**). The disruption of B-form DNA is reinforced by the equivalent residue in MCM2, W569, that engages in a similar interaction, by stacking against the base of the 5′-end thymidine (**Figure 3**). All MCMs, with the exception of MCM3, have either an aromatic or a bulky hydrophobic residue at this position, suggesting that they can all in principle engage in a similar interaction (**Supplementary figure 7**). The position of MCM6 F442 at the MCM-DNA interface is reminiscent of that of H507 in the DNA-binding loop of the viral E1 helicase (Enemark & Joshua-Tor, 2006). As proposed for E1 H507 (Liu, Schuck, & Stenlund, 2007), the aromatic/hydrophobic residue of the [+]x_6_[Ω/Ψ] motif might have an additional or alternative role at an earlier stage in replication, by helping drive melting of origin DNA.

Correct positioning of the invariant serine at the start of the H2I loop for interaction with the phosphate backbone of the DNA requires adoption of a helical conformation by the five residues succeeding the serine (H2I *α*_N_; **Figure 3** and **Supplementary figure 7**). This local helical folding is driven by anti-parallel β-strand pairing of the two residues preceding the serine with the second β-strand of the PS1 loop in the preceding MCM subunit. This inter-subunit interaction helps merge the H2I and PS1 loops of individual MCMs into a continuous DNA-binding staircase that extends around the pore in the C-tier, as noted recently for the archaeal homo-hexameric MCM ring (Meagher, Epling, & Enemark, 2019). As expected for the MCM subunit at the bottom of the staircase, H2I *α*_N_ is disordered in MCM5.

### ATP binding and hydrolysis

In the structure, five of the six ATP-binding sites in the MCM2-7 ring are occupied by a nucleotide (**Figure 4** and **Supplementary figure 8B, C**). The three contiguous ATP-binding sites of MCM6, MCM4 and MCM7 contain ATPγS, with a Mg^2+^ ion coordinated between β and γ phosphates. Interestingly, the cryoEM map shows clearly that MCM2 and MCM3 have hydrolysed their ATPγS to ADP. Furthermore, the MCM5 nucleotide-binding site is empty. It is possible that MCM5 had hydrolysed ATPγS and released ADP, or alternatively that ATPγS was never bound: both possibilities are compatible with the observed open state of the MCM5-2 interface.

**Figure 4.**
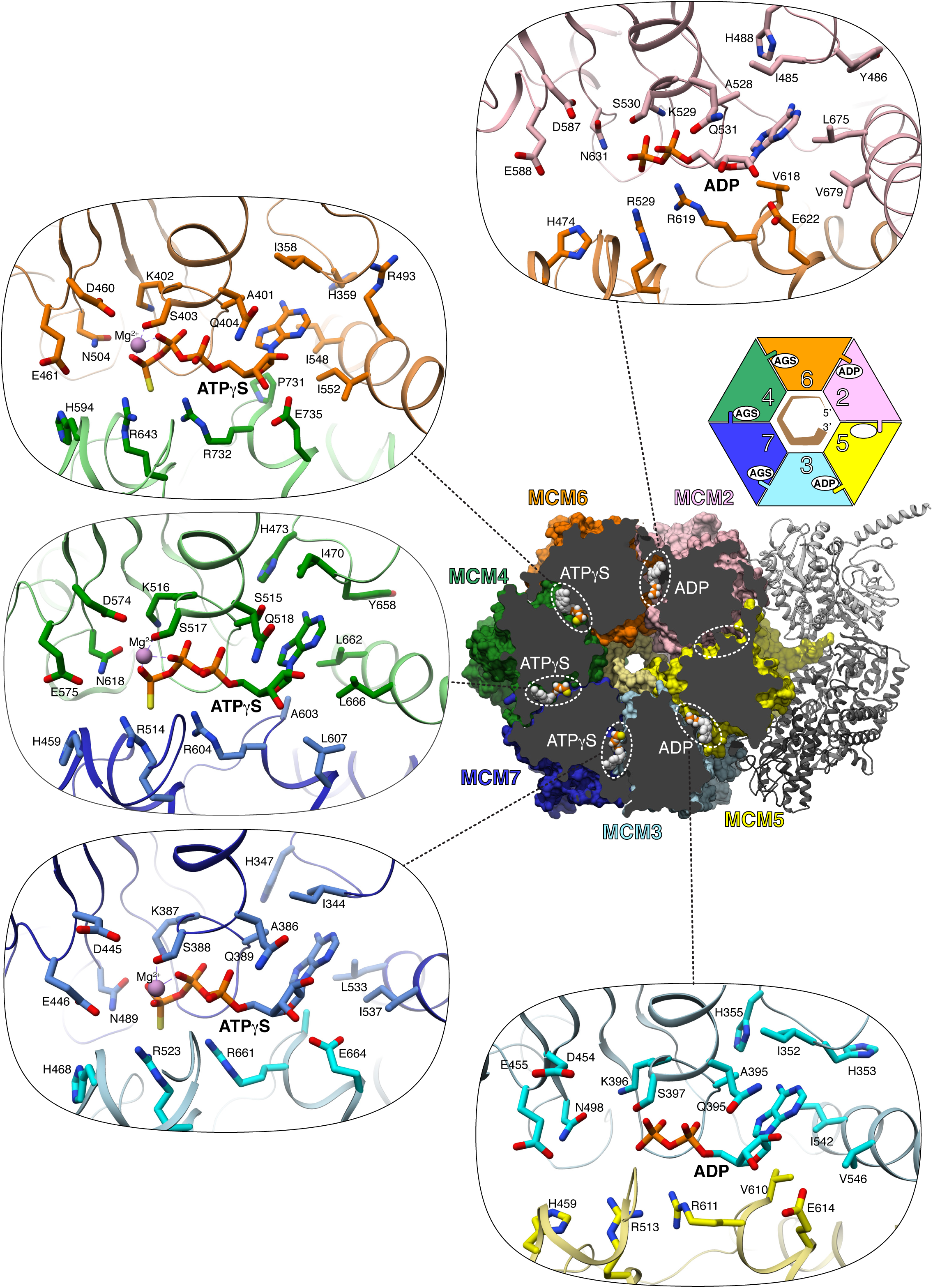
ATPγS binding in the CMG-ATPγS-DNA complex. In the middle-right panel, the MCM C-tier is shown in molecular surface representation, clipped from the N-tier end, to reveal the nucleotide bound at each MCM interface. The cartoon drawing above the panel recapitulates the ATP status of MCM2-7 (AGS stands for ATPγS). Details of nucleotide binding are provided in the oval panels for each of the five nucleotide-bound interfaces. MCM chains are colour-coded as in Figure 1.

All residues previously identified as involved in ATP binding and hydrolysis engage as expected with the ATPγS moieties bound at the three MCM7-3, MCM4-7 and MCM6-4 interfaces, including Walker A and B residues and sensor-1 asparagine of the P-loop subunit, and arginine finger, sensor-2 arginine and sensor-3 histidine residues in the contiguous ‘sensor’ subunit (**Figure 4** and **Supplementary figure 9**). The two ADP-bound interfaces between MCM3-5 and MCM2-6 show a very similar set of contacts, except that the arginine fingers in MCM5, R513, and MCM6, R529, are partially disordered, likely as a consequence of the absence of the γ phosphate. A noteworthy feature of ATP binding by MCM2-7 is the extensive range of hydrophobic interactions that shield the aromatic base of the nucleotide from solvent (**Figure 4**). These interactions include a ‘sandwich’ interaction made by an invariant isoleucine on one side of the base and two aliphatic residues in the domain-swapped helix on the other side (**Figure 4**). In addition, the adenine base engages in Watson-Crick like hydrogen bonding with the main-chain nitrogen and carbonyl moieties of a residue preceding a conserved ‘positive Φ’ glycine in the linker between helices 2 and 3 of the ATPase domain (**Supplementary figure 10**).

An unresolved question is whether ATP-coupled conformational changes during DNA translocation are limited to movements of the DNA-binding loops or involve the entire ATPase domain. Structural superposition of the ATPase domains shows that the DNA-binding loops of nucleotide-bound MCM6, 4, 7 and 3 are in similar position relative to their ATPase domains (**Figure 5**); in contrast, the H2I loop of apo MCM5 at the 3′-end of the DNA occupies a lower position and its H2I *α*_N_ is disordered (**Figure 5**). These observations suggest that DNA translocation might be achieved by a composite mechanism of whole-domain movements during the ATP-coupled translocation cycle, and rearrangement of the H2I loop at the end of the cycle, as H2I detaches itself from the staircase and its ATPase domain re-engages DNA at the 5′-end.

**Figure 5.**
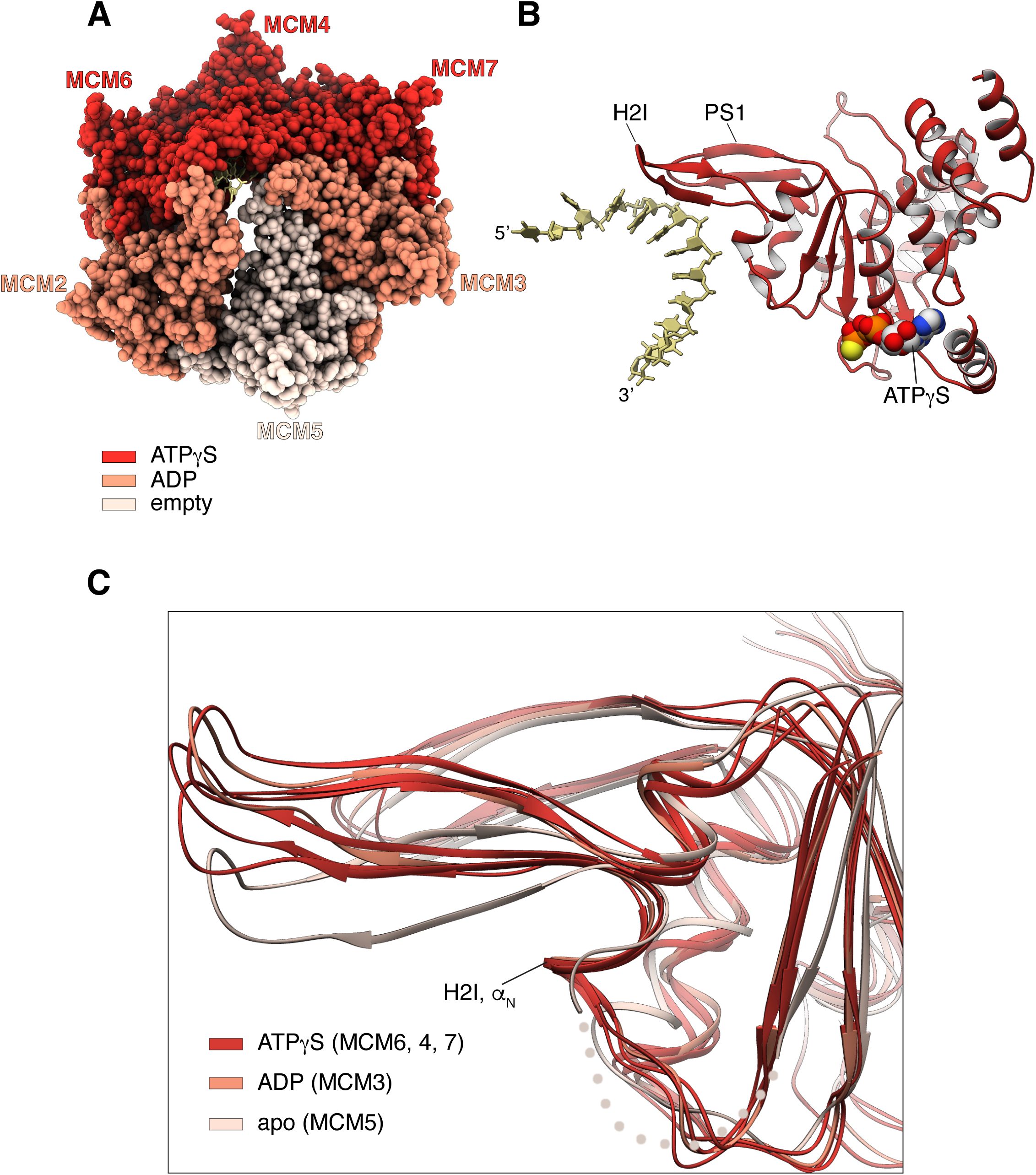
Conformational coupling of ATP status and DNA-binding loops. **A** Spacefill representation of the MCM C-tier, colour-coded according to ATP status. **B** Ribbon representation of MCM4 and DNA, showing the reciprocal position of the ATPγS moiety and the H2I and PS1 DNA-binding loops. **C** Superposition of MCM ATPase domains, highlighting the relative position of their DNA-binding loops (MCM2 is omitted, see Supplementary figure 11). The H2I *α*_N_ helix of MCM5 is disordered and drawn as a dotted line. The MCM ATPase domains are coloured according to ATP status, as in panel A.

The ATP status of the ADP-bound MCM2 subunit appears anomalous given its position at the top of the ring staircase. Several indicators point to MCM2 acting as a ‘seam subunit’ (Eickhoff et al., 2019) that has only partially engaged with the rest of the C-tier ring: the smaller interface area with MCM6 (1391 Å^2^, instead of ∼2000 Å^2^ for the other nucleotide-occupied MCM interfaces), the disordered conformation of its DNA-binding element H2I *α*_N_, and the higher B value attained during real-space refinement. The ADP moiety of MCM2 is also unusual in the way it sits in the P-loop, as the nucleotide is shifted so that its β phosphate occupies the position occupied by the γ-phosphate in the ATPγS-bound interfaces (**Supplementary figure 11**).

Overall, the observations relative to ATP status and DNA binding in the MCM C-tier ring are consistent with a sequential rotary mechanism of ATP hydrolysis as the basis for translocation of the human CMG. The structural integrity of the C-tier during translocation is provided by a domain-swapped helix (**Supplementary figure 4**) that provides a flexible tether between contiguous MCM subunits, in a similar fashion as recently described for the replicative gp4 DNA helicase (Gao et al., 2019).

### Interaction with AND-1

Analysis of purified human CMG overexpressed in HEK293 cells revealed the co-purification of sub-stoichiometric amounts of endogenous AND-1, a known replisome factor and the human orthologue of yeast Ctf4 (**Supplementary figure 1**). Our previous work had shown that yeast Ctf4 acts as a recruitment hub for replisome proteins, tethering multiple factors at the fork via its trimeric structure (Simon et al., 2014; Villa et al., 2016). AND-1 shares its oligomeric nature with Ctf4, although it appears to have a distinct mechanism of binding to its replisome partner, Pol *α*/primase (Kilkenny et al., 2017).

Co-purification of endogenous AND-1 indicated a strong constitutive interaction with the human CMG complex. We therefore decided to co-express AND-1 together with the components of the human CMG, and succeeded in purifying a 12-subunit CMG assembly which we refer to here as CMGA (**Supplementary figure 12**). CryoEM analysis of CMGA using Warp for particle picking (Tegunov & Cramer, 2019) and CryoSparc for image reconstruction (Punjani, Rubinstein, Fleet, & Brubaker, 2017) yielded a 6.77 Å map that was readily interpretable and permitted the unambiguous docking of the high-resolution structure of human CMG and the crystal structure of the AND-1 trimer that we reported earlier (Kilkenny et al., 2017) (**Supplementary figure 13**).

The structure shows that the disk-shaped AND-1 trimer docks edge-on at a near perpendicular angle onto the leading face of the CMG (**Figure 6**). Despite AND-1 being full-length, only the SepB-like domain of AND-1 is visible in the map, indicating that the N-terminal β-propeller domain and its extended C-terminal portion spanning the HMG box are flexibly oriented in the trimeric structure. Relative to the CMG, the trimeric AND-1 disk is arranged so that its N-terminal segments are located ahead of the fork and in proximity of the parental double-stranded DNA. In contrast, the helical structure of the SepB domain and the relative C-terminal extensions project away from the CMG (**Figure 6**).

**Figure 6.**
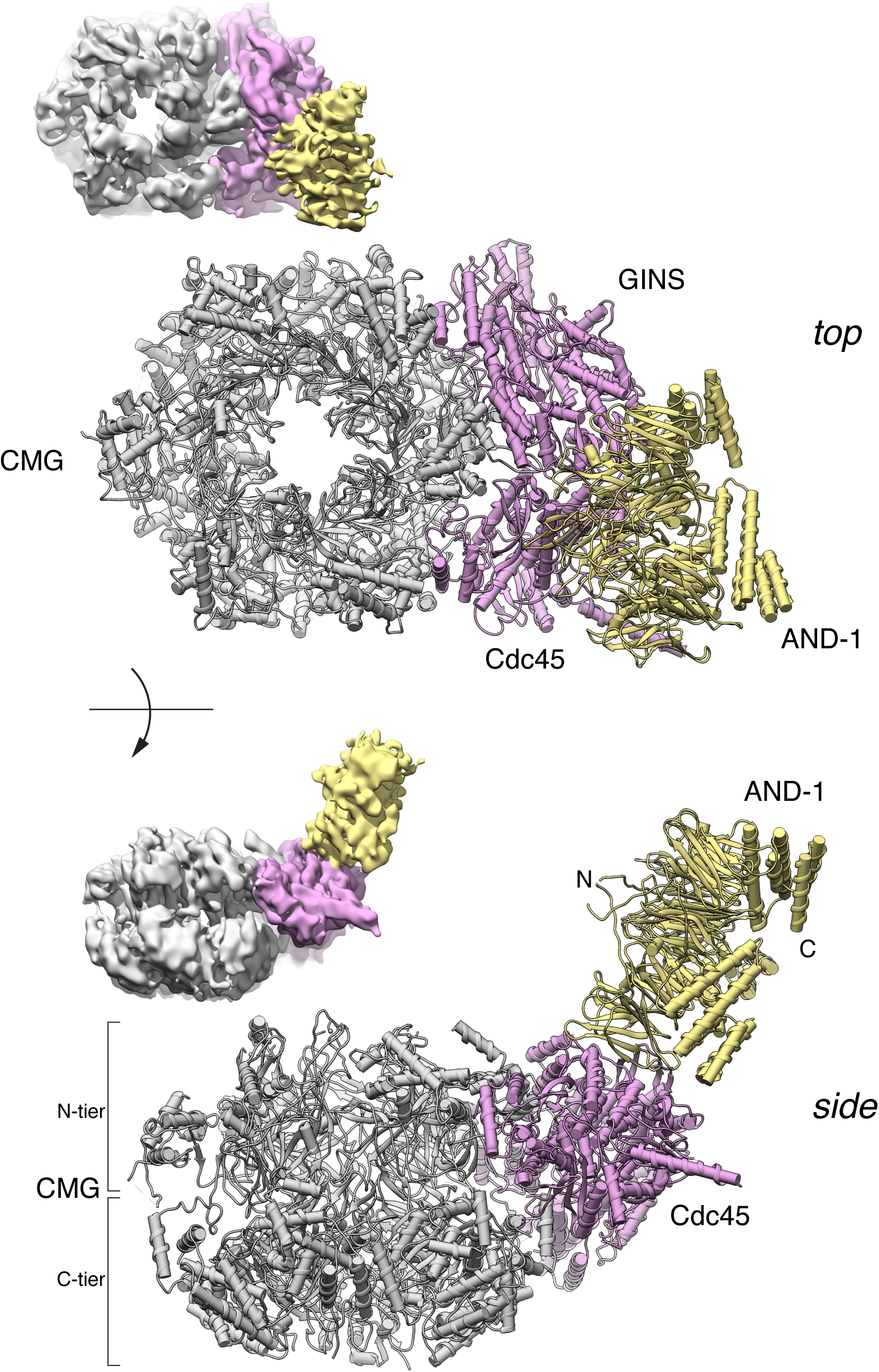
Structure of human CMG bound to AND-1 (CMGA). Top and side views of the atomic model of CMGA are shown in ribbon drawing. The CMG is coloured in grey, Cdc45 and GINS in pink and AND-1 in yellow. At the top-left of each ribbon drawing is reported a view of the cryoEM map in the same orientation, coloured accordingly.

AND-1 binds at the perimeter of the CMG, engaging both Cdc45 and GINS with the first and last blade of one of its β-propeller domains (**Figure 7**). The CMG - AND-1 interface is formed by the B-domain of Psf2 that projects towards the concave surface formed by blades 1 and 6 of AND-1’s β-propeller, as well as the helical portion of Cdc45 that links its two DHH domains. The interface buries only 1087 Å^2^, a surprisingly small area for a constitutive interaction (**Figure 7**). The limited resolution of our structure is insufficient for unambiguous identification of interface amino acids. However, we can determine that the interface is of mixed hydrophobic and hydrophilic nature, and that the tight binding of AND-1 to the CMG despite the relatively limited interface might be driven by the presence of charge-charge interactions that become solvent-excluded upon CMGA formation.

**Figure 7.**
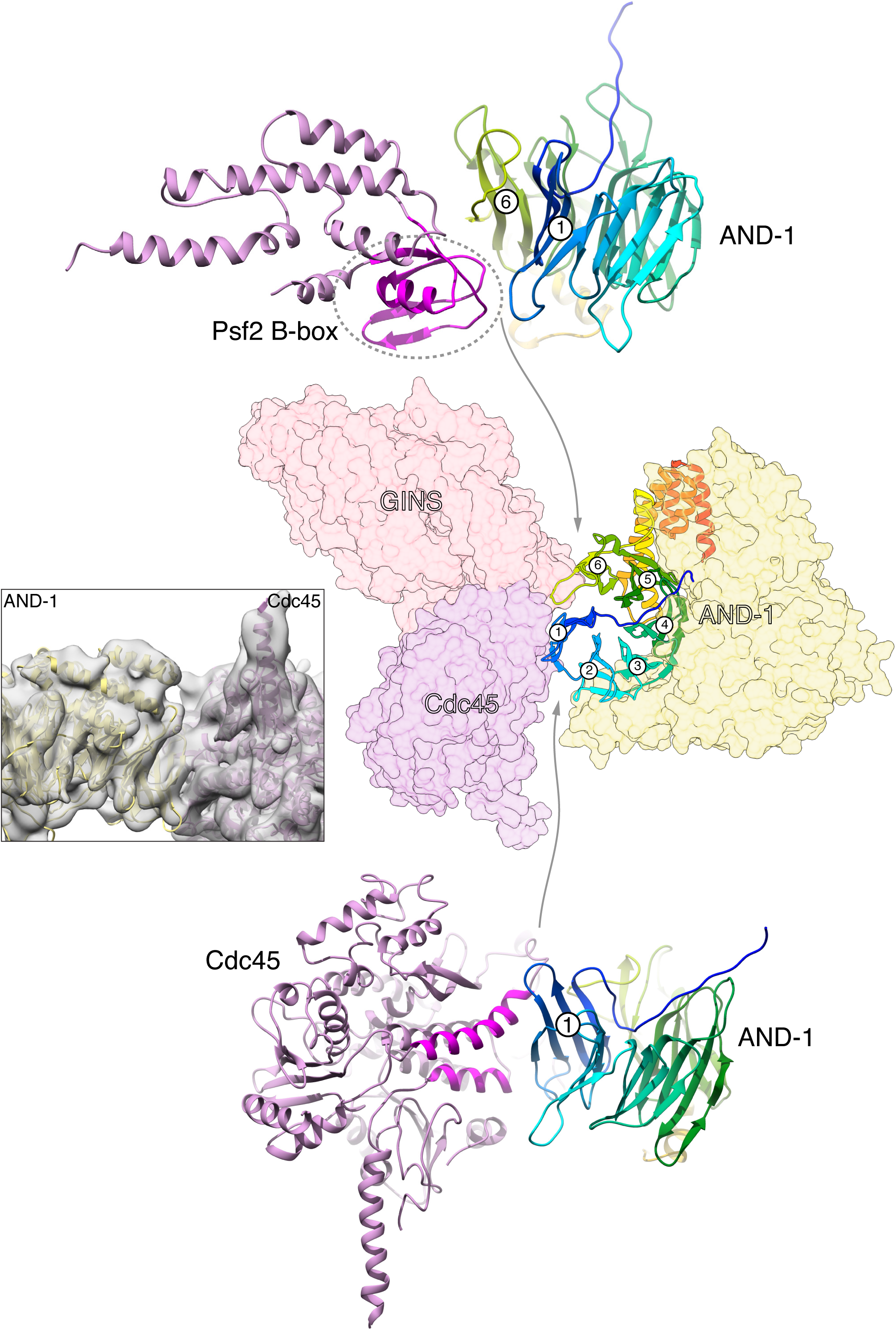
The CMG - AND-1 interface in CMGA. The middle panel shows GINS (light pink), Cdc45 (darker pink) and AND-1 (yellow) as transparent molecular surfaces. The β-propeller domain of the AND-1 trimer that contacts the CMG is drawn as ribbon, coloured from blue (N-end) to red (C-end). The six blades of the propeller are numbered 1 (N-end) to 6 (C-end). To the left is a view of the cryoEM map for the CMGA, with ribbon models of AND-1 (yellow) and Cdc45 (pink) fitted in the map. Above and below the central panels are ribbon drawings showing the interface of AND-1’s β-propeller domain with the B-box of Psf2 (top) and with Cdc45 (bottom).

## DISCUSSION

In this paper, we have used cryoEM to capture a high-resolution view of the human CMG bound to a fork DNA substrate in the presence of ATPγS. Our map of human CMG-ATPγS-ssDNA allowed us to visualise unambiguously critical features of the complex, such as its protein-DNA interface and the nucleotide-binding sites, and to represent them in an accurate atomic model. We have also reported an intermediate-resolution structure of the CMGA assembly, which described the mode of interaction of CMG with the core replisome factor AND-1.

### DNA binding

In our structure, all six MCM subunits contact ssDNA, spanning a total of 11 nucleotides. The footprint of an MCM subunit on ssDNA covers four nucleotides, rather than two as previously reported (**Figure 3**) (Eickhoff et al., 2019), with two overlapping nucleotides between neighbouring subunits. Interaction of MCM6, 4, 7 and 3 with DNA takes place via an identical set of contacts mediated by both PS1 and H2I DNA-binding loops (**Figure 3**). A significant difference is that MCM6 and MCM2 use aromatic residues at a conserved H2I position to unstack the nucleotides at the 5′-end of the DNA and disrupt the B-form DNA. These aromatic residues intermesh with consecutive bases much as the teeth of a cogwheel; such contacts appear well suited to avoid slippage and transmit torque when an MCM subunit engages the leading DNA strand emerging from the N-tier, at the top of the staircase. Overall, the arrangement of PS1 and H2I loops within the C-tier ring follows closely the DNA spiral, lowering steadily their vertical reach from MCM2 at the top of the binding staircase to MCM5 at the bottom (**Figure 2**).

Earlier structural work on yeast and fly CMG had shown that DNA can be bound via two different sets of MCM subunits: MCM6, 4 and 7 (Abid Ali et al., 2016) or MCM2, 3, 5 and 6 (Georgescu et al., 2017; Goswami et al., 2018). A recent cryoEM analysis of fly CMG in the act of translocating on DNA revealed the existence of several different conformational states, which appear to encompass and extend the previously described DNA-bound states of the CMG (Eickhoff et al., 2019). In light of this analysis, our CMG-DNA structure would most likely correspond to state 2B, in which MCM6, 4, 7 and 3 contact DNA and are bound to ATP, with MCM2 in the process of exchanging ADP for ATP and re-engaging with DNA at the 5′-end. Thus, the emerging evidence from this wealth of structural data strongly indicates multiple modes of asymmetric DNA binding in the C-tier ring as a key feature of DNA translocation by the CMG.

### ATP site occupancy and hydrolysis

The site occupancy and hydrolysis status of ATPγS in our CMG-DNA structure is in general agreement with the sequential rotary model of ATP utilisation but also reveals some unexpected findings. Nucleotide occupancy is known to correlate with DNA binding, and in our structure ATPγS-bound MCM6, 4 and 7 interact with DNA. In the model, ring subunits at or near the bottom of the DNA-binding staircase have hydrolysed ATP; accordingly, the MCM3-5 interface has converted the slowly-hydrolysable ATPγS to ADP (**Supplementary figure 8C**), whereas MCM5 is in the apo state. These observations are in line with the sequential rotary model, however they represent an apparent discrepancy with the model of Eickhoff and colleagues (Eickhoff et al., 2019), in which ATP binding by MCM3, but not its hydrolysis, is important for asymmetric translocation mediated by a MCM3-5 dimer. Interestingly though, the ADP-bound MCM3-5 interface remains as extensive as for the three ATPγS interfaces of MCM6-4, MCM4-7 and MCM7-3 (2078 Å^2^ of buried surface area versus an average value of 2034 Å^2^ for the ATPγS interfaces).

Furthermore, in our structure the MCM5-2 interface is void of nucleotide and the MCM2-5 gate is open. Given the finding that MCM3 has hydrolysed ATPγS, MCM5 might be one step ahead in the ATP cycle and may have released its ADP, in preparation for re-joining the C-tier ring at the top of the staircase. That MCM3 and MCM5 of all six subunits should have hydrolysed ATPγS is in agreement with the effect of Walker A K-to-A mutations in fly CMG, showing that loss of ATP binding by MCM3 and MCM5 caused the largest decrease in ATP hydrolysis rates (Ilves et al., 2010), and with a four-fold reduction in DNA unwinding caused by an ‘arginine finger’-to-alanine mutation in fly MCM5 (Eickhoff et al., 2019). Whether the open state of the apo MCM2-5 interface represents a natural intermediate state in the translocation cycle, or rather a stalled or paused state of the helicase remains to be established. At any rate, the unexpected indication of ATPγS hydrolysis provides evidence that the CMG has engaged productively with the fork DNA substrate and might have undergone a limited degree of translocation.

The ADP-bound status of MCM2, at the top of the staircase and interacting with the 5′-end of the ssDNA, is apparently inconsistent with a sequential rotary model of ATP hydrolysis. We have already described several elements of evidence indicating that the MCM2 appears to behave as a ‘seam’ subunit (Eickhoff et al., 2019). An intriguing possibility is that the observed ATPγS hydrolysis by MCM2 might be a clue of translocation with reverse polarity, that could have been prompted by idling of the helicase upon incubation with slowly-hydrolysable ATPγS. Evidence that the CMG can backtrack on ssDNA has been provided by recent single-molecule studies (Burnham, Kose, Hoyle, & Yardimci, 2019).

### DNA translocation

The staircasing arrangement of the DNA-binding loops and the ATP-hydrolysis status in the MCM ring are both supportive of a sequential rotary mechanism of DNA translocation for the human CMG. Differences, such as the observed ATP-hydrolysis status of MCM3, with the proposed model of asymmetric translocation (Eickhoff et al., 2019), remain to be explained and might be species-specific.

Our structure captures a high-resolution snapshot of the CMG trapped on fork DNA with ATPγS and does not provide conclusive evidence concerning models of asymmetric translocation. However, insight into asymmetry in MCM2-7 behaviour comes from analysis of the solvation free-energy for formation of the interfaces between contiguous ATPase domains (Δ^i^G) in the C-tier ring, using the EBI PISA server (Krissinel & Henrick, 2007). The analysis shows striking differences in Δ^i^G values among MCM interfaces. Although the three ATPγS interfaces, as well as the ADP-bound MCM3-5 interface, bury similar surface areas (between 1975 Å^2^ and 2140 Å^2^), formation of the MCM4-7_ATPγS_ and MCM6-4_ATPγS_ interfaces shows more than 2-fold higher energy gains and 7-fold lower P-values than for the MCM7-3_ATPγS_ and the MCM3-5_ADP_ interfaces (**Supplementary figure 14**). Thus, the Δ^i^G analysis indicates that the contacts binding together the MCM 4-7 and 6-4 interfaces are much tighter and more specific than in the other MCM interfaces. These observations might provide a structural basis for the finding that human MCM4, 6 and 7 can be recovered as a stable heterotrimeric complex from HeLa cells (Ishimi, 1997). They further suggest that MCM subunits 4, 6 and 7 might behave as a rigid body in the asymmetric mode of DNA translocation. This would be in agreement with biochemical evidence that loss-of-ATP-binding K-to-A mutations in fly MCM6 and 4 causes only modest reductions in DNA unwinding (Ilves et al., 2010), and that loss of ATP hydrolysis at the MCM6-4 and MCM4-7 interfaces caused by alanine mutation of the ‘arginine finger’ in MCM4 and 7 has equally modest effects on unwinding (Eickhoff et al., 2019).

The evolutionary invariance of catalytic residues in all six MCM2-7 ATPases represents a challenge for models of asymmetric translocation that consider ATP binding and hydrolysis to be important for a subset of MCM subunits. The following points in this regard can be made: as far as we can determine in our cryoEM map, all catalytic and sensor MCM residues at each nucleotide-bound interface engage correctly with the nucleotide and are therefore potentially capable of catalysis. Furthermore, symmetric translocation is adequate for DNA replication by homo-hexameric DNA helicases in viruses, bacteria and archaea, implying that it represents the evolutionary consensus for DNA translocation. Finally, differentiation of the eukaryotic MCM into six distinct proteins may have evolved to endow the CMG with its unique mode of loading and activation and possibly termination. Asymmetric translocation might therefore represent an adaptation to cope with MCM sequence diversification rather than a process of optimisation. Consequently, the eukaryotic CMG might be capable of considerable mechanistic flexibility, concerning the role played by each of its MCM subunits during translocation.

### CMGA

Our cryoEM structure of the human CMGA elucidates the interaction mechanism of the replisome component AND-1 with the CMG. AND-1 does not contact the MCM proteins and binds to the side of the CMG where Cdc45 and GINS are located. Thus, an important role of Cdc45 and GINS, in addition to activating the CMG helicase activity, is to mediate CMG’s interaction with AND-1. Only the trimer of AND-1’s SepB-like domains is visible in the map, indicating that its N-terminal β-propeller and extended C-terminal region are flexibly arranged relative to its central trimeric structure. AND-1 is docked onto the CMG like a rigid body, with a single β-propeller wedged in between Cdc45 and GINS. This mode of CMG binding is similar to the mode of interaction that was recently described for yeast Ctf4 with the CMG (Yuan et al., 2019), suggesting that the resulting architecture of the CMGA is functionally important and has been evolutionarily preserved.

Because of the high interaction angle of the disk-like AND-1 trimer relative to the plane containing the leading edge of the CMG, the N-terminal β-propeller domains of AND-1, not visible in our structure, would be placed in the trajectory of the parental DNA. This striking feature of the CMGA architecture indicates that AND-1/Ctf4 can in principle contact the parental DNA ahead of the CMG’s leading edge of translocation. Clearly, these findings point to further discoveries and unexpected observations that await future structural studies of the eukaryotic replisome.

We had originally reported that Ctf4 can interact with the CMG via a CIP (Ctf4-Interacting Peptide) motif present in the N-tail of yeast Sld5 (Simon et al., 2014), as well as in Ctf4’s multiple protein partners (Villa et al., 2016). In the light of our current observations, we believe that the mode of AND-1 interaction with the CMG described here and reported for yeast Ctf4 represents the principal mode of AND-1/Ctf4 recruitment to the replisome. Binding by the CIP of yeast Sld5 might further secure the association of Ctf4 to the yeast CMG, as well as possibly act as a safety mechanism for keeping Ctf4 anchored to fork when its primary interaction site with the CMG is disrupted. A CMG-binding motif equivalent to the yeast CIP is not found in human Sld5 or any other human CMG components, indicating that the Ctf4-Sld5 CIP interaction is unique to budding yeast.

## METHODS

### Construct design and preparation

For expression of human CMG, ORFs for full-length human MCM2 (P49736), MCM3 (EAX04367), MCM4 (P33991), MCM5 (P33992), MCM6 (Q14566), MCM7 (P33993), Cdc45 (AAC67521), Psf1 (Q14691), Psf2 (Q9Y248), Psf3 (Q9BRX5) and Sld5 (Q9BRT9) were synthesised using the GeneArt Gene Synthesis service (ThermoFisher). GenBank EAX04367 codes for an MCM3 isoform (853 aa) that contains 45 additional residues at the N-terminus relative to Swissprot P25205.

ORFs were codon optimised for overexpression in human cells and designed with flanking restriction sites for insertion into the ACEMam1 and 2 vectors of the MultiMam transient system (Vijayachandran et al., 2011). MCM4 was encoded with an N-terminal His_8_-TEV tag, while Psf2 was encoded with a C-terminal TEV-2xStrepII tag. Multi-cassette constructs encoding MCM4-6-7, MCM2-3-5 and Cdc45-Psf1-Psf2-Psf3-Sld5 were generated making use of the I-CeuI / BstXI sites in the MultiMam vectors. For expression of human CMG - AND-1 (CMGA), the full-length ORF of human AND-1 (IMAGE cDNA clone 6514641) was cloned with an N-terminal 2xStrepII-TEV tag into the ACEMam2 construct expressing MCM2-3-5, while the C-terminal Strep tag fused to the Psf2 gene was removed.

### Mammalian tissue culture and protein production

HEK293-F cells (FreeStyle™, ThermoFisher #R79007) were maintained in suspension culture according to manufacturer’s instructions, using Gibco serum-free FreeStyle™ Media (#12338018). Cultures were kept at 37 °C, 8% CO_2_ and 70% humidity in a Kuhner ISF1-X incubator shaking at 125 rpm. For large-scale transient transfection, HEK293-F cells were grown in 400 mL batches in 2 L non-baffled Erlenmeyer flasks (TriForest Enterprises Inc, #FPC2000S) to a cell density of ∼1×10^6^ cells/mL.

For each flask, a separate transfection mixture was prepared as follows: a total of 480 μg recombinant plasmid DNA encoding CMG or CMGA was added to 40 mL fresh media. Equimolar ratios of the three relevant recombinant plasmids were used. 960 μg PEI (Polyethylenimine, linear, MW∼25,000; Polysciences Inc, #23966) was added from a sterile 1 mg/ml stock prepared in 50 mM Hepes-KOH pH 7.5. The resulting transfection mixture was vortexed vigorously for 10 seconds, then incubated at room temperature for 15 minutes before being decanted into the cell culture.

Three hours post-transfection, 4 mM valproic acid (Sigma, #P4543) was added to the cultures, to enhance protein expression. Cultures were returned to shaking incubation for four days, before being harvested by centrifugation at 500 x g for 10 minutes at 10 °C. Cell pellets from ∼1.2 L cell culture were subsequently resuspended in 40 mL of chilled, sterile PBS supplemented with SIGMAFAST EDTA-free protease inhibitor cocktail (Sigma, #S8830). Washed cells were again harvested by centrifugation, then snap-frozen in 50 mL centrifuge tubes using liquid nitrogen, and stored at -80 °C.

### Protein purification

To purify human CMG, a frozen cell pellet from ∼1.2 L cell culture was thawed on ice in 35 mL Buffer N (50 mM Hepes-KOH pH 7.5, 5% v/v glycerol, 300 mM potassium glutamate, 5 mM magnesium acetate) supplemented with 40 mM imidazole (pH 7.5), 0.02% v/v IGEPAL (Sigma, #I3021), 5 μL Benzonase (Sigma, #E1014), 0.5 mg bovine pancreas RNaseA (Sigma, #R6513), and SIGMAFAST EDTA-free protease inhibitor cocktail (Sigma, #S8830). The resulting suspension was subsequently sonicated on ice for a total of 2 minutes (5-second bursts with 10-second recovery) using a 6 mm probe and a Sonics Vibra-Cell VCX 130 set to 60% amplitude.

Cell lysate was clarified by centrifugation at 45,000 x g for 1 hour at 4 °C, then filtered through a 5 μm Acrodisc syringe filter (PALL, #4650), a 0.45 μm HV Durapore vacuum filter (Millipore, #SCHVU01RE) and finally a 0.2 μm Minisart syringe filter (Sartorius, #16534-K). The filtered sample was applied to a 5 mL HisTrap HP column (GE Healthcare, #17-5248-02), pre-equilibrated with Buffer N and 40 mM imidazole using a peristaltic pump. All subsequent chromatography steps were performed on an ÄKTA Purifier (GE Healthcare). The loaded column was washed twice with 5 column volumes of Buffer N with 40 mM imidazole and bound protein was eluted in reverse flow using Buffer N with 300 mM imidazole. 2 mL fractions corresponding to ∼10 mL of the elution peak were pooled and applied to a 1 mL StrepTrap column (GE Healthcare, #28-9075-46) pre-equilibrated with Buffer N with 2 mM DTT. After sample loading, the column was washed with 10 column volumes of Buffer N and 4 column volumes of Buffer S (25 mM Hepes pH 7.5, 50 mM potassium chloride, 5 mM magnesium acetate, 2 mM DTT). The CMG was eluted in reverse flow using Buffer S with 15 mM d-Desthiobiotin (Sigma, #D1411); 0.5 mL fractions corresponding to the ∼1.5 mL elution peak were analysed by SDS-PAGE and stored at 4 °C.

To purify the CMG-DNA complex, the CMG purification protocol was followed until the StrepTrap column loading step. After sample loading, the column was washed for 5 column volumes using Buffer N with 2 mM DTT, followed by 5 column volumes of Buffer S. To form the CMG-DNA complex, ∼900 μL of 4 μM forked DNA duplex and 100 μM ATPγS (Jena Bioscience, #NU-406-50) in Buffer S was applied to the column at 0.05 mL/min. The column was first washed in reverse flow for 4 column volumes with Buffer S and 100 μM ATPγS, and the CMG-DNA complex was eluted in reverse flow using Buffer S, 100 μM ATPγS and 15 mM d-Desthiobiotin. 0.5 mL fractions corresponding to the ∼1.5 mL elution peak were analysed by SDS-PAGE and stored overnight at 4 °C. The peak fraction was used for grid preparation.

To purify the CMGA complex, the CMG purification protocol was followed using cell cultures that had been transfected with the ACEMam2 construct expressing AND-1 and MCM2-3-5.

### Preparation of fork DNA

A fork DNA substrate was generated using two 70 bp oligonucleotides, based on a design by Petojevic and colleagues (Petojevic et al., 2015): CGTTTTACAACGTCGTGACTGGGCACTTGATCGGCCAACCTTTTTTTTTTTTTTTTT TTTTTTTTTTTTT (70T) CAGGCAGGCAGGCAGGCAGGCAGGCAGGCAGGTTGGCCGATCAAGTGCCCAGTCACGACGTTGTAAAACG (70GC)

Oligonucleotides were supplied PAGE-purified by IDT and resuspended to 100 μM in TE buffer pH 8 (Invitrogen, #AM9849). Equimolar amounts were mixed in 200 μL aliquots at 40μM in TE supplemented with 50 mM NaCl, prior to annealing. The resulting fork DNA consisted of a 40 bp duplex region and a 30 nt fork comprising a 3*′* poly-dT tail for CMG loading in the correct orientation for unwinding and a 5*′* GC-rich tail that does not bind CMG (Petojevic et al., 2015).

### CryoEM sample preparation and data collection

UltrAuFoil R1.2/R1.3 300 mesh gold grids (Quantifoil Micro Tools GmbH, #N1-A14nAu30-01) were glow-discharged twice (once on each side) for 1 minute using a PELCO easiGlow system (0.4 mBar, 30 mA, negative polarity). Grid samples were then prepared using a Vitrobot Mark IV robot (FEI), set to 100% humidity, 4 °C, 2.5 seconds blot time and -10 blot force. 3 μL of CMG-DNA complex was applied to both sides of the grid prior to vitrification in liquid ethane.

Cryo-EM grids of CMGA complex were prepared as above, with two variations. Firstly, ATPγS was added to the protein sample at a final concentration of 1 mM approximately 1 hour before grid preparation. Secondly, a 5% glutaraldehyde cross-linking solution was added in 1:10 volume ratio to the protein sample approximately 10 minutes before grid preparation. The protein sample was incubated on ice at all times.

Grid samples were initially screened on a Talos Arctica (FEI) operating at 200 keV. High-resolution data were subsequently acquired for a single grid on a Titan Krios (FEI) operating at 300 keV. Automated data collection for Single Particle Analysis (SPA) was performed using the EPU package (FEI). Grid preparation, screening and data collection were performed at the Cryo-EM facility in the Department of Biochemistry.

### CryoEM data processing

Statistics for data collection and processing are reported in Table S1.

#### CMG-DNA

Data processing was performed on the Cambridge Service for Data-Driven Discovery (CSD3) high-performance computer cluster, using RELION-3 (Zivanov et al., 2018). Motion correction for all 3,694 movies was performed using 5 x 5 patch alignment in MotionCor2 (Zheng et al., 2017) with ‘InFmMotion’ activated to take into account frame motion blurring. CTF correction was performed using GCTF (Zhang, 2016) against non-dose weighted averages, with ‘equiphase averaging’ activated. Micrographs yielding resolution limit estimates of >6 Å were discarded. Laplacian-of-Gaussian-(LoG-) based auto-picking with a default threshold of -0.1 was used to pick a total of 845,596 particles across 3,619 micrographs. This particle set was used for all downstream processing without the use of template-driven picking procedures. Particles were initially extracted 4xbinned with a box size of 90 pixels (4.28 Å/pixel). For iterative rounds of 2D classification, the option to ‘Ignore CTFs until first peak’ was activated. This yielded a final set of 365,202 high-resolution CMG particles, which was then subjected to 3D classification using an internally generated initial model. Particles from 3 out of 4 resulting classes were pooled (297,183 total), re-extracted without binning (360 pixels, 1.07 Å/pixel) and refined against a suitably re-scaled initial model. Subsequent soft mask generation and re-refinement with solvent flattened FSCs yielded a first high-resolution map of CMG at 3.50 Å, with clear density for ssDNA in the pore. CTF refinement, Bayesian particle polishing and additional masked refinement of the polished particles using solvent flattened FSCs improved the resolution to 3.24 Å.

Inspection of the polished map identified some unresolved heterogeneity in the structure, particularly in the C-tier ATPase domains of the MCM ring. The unbinned polished particle set was therefore subjected to a further round of 3D classification (3 classes) without alignments. The majority of the particles (213,527) formed a single DNA-bound CMG class, which was used for further processing. A second DNA-bound CMG class (30,696 particles) was deemed poor quality and discarded, while a third CMG class (52,960 particles) did not have bound DNA.

The high-resolution DNA-bound CMG class was subjected to refinement with solvent flattened FSCs, yielding a final resolution of 3.29 Å and providing clearer definition of the C-terminal ATPase domains. This model was used for the majority of model building. However, ATPase domains for MCM2 and 5 still exhibited some heterogeneity. To resolve this, all particles were first shifted to the centre of mass of the MCM2-7 ATPase ring and density outside this ring was subtracted from the shifted particles, facilitating focussed 3D classification (without alignment) of the MCM2-7 C-tier. The majority of particles (181401, 85%) populated the first of two classes. This presented improved definition for the ATPase domains of MCM2 and 5, and also revealed good density for the C-terminal domains of MCM2 and 6. Masked, solvent-flattened refinement of this class resulted in a C-tier map extending to 3.41 Å that was used to complete model building.

#### CMGA

Data processing was performed using Warp (Tegunov & Cramer, 2019) for motion correction, CTF estimation, automated particle picking and particle extraction, and cryoSPARC (Punjani et al., 2017) for all subsequent steps. A total of 140,128 particles were automatically picked and extracted from 1,844 micrographs. An initial round of 2D classification identified 6 classes that presented sensible densities; contributing particles were used to generate two distinct initial models that were characterised as CMG (18,439 particles) and AND-1 (9,600 particles). The CMG initial model exhibited weak additional density adjacent to the GINS/Cdc45 subunits, suggesting a mixed population of CMG and CMGA in the data. Accordingly, the CMG initial model and particle set were subjected to hetero-refinement for 2 classes, resulting in a CMG-only particle set (7,860 particles) and a CMGA particle set (10,579 particles). The latter was again subjected to hetero-refinement for 2 classes to remove any remaining CMG-only particles, resulting in an improved CMGA map derived from 8,888 particles.

The highest quality CMG, CMGA and AND-1 models derived from the initial 2D classification were then used to drive iterative rounds of hetero-refinement against the entire 140,128-particle dataset. Once suspected AND-1 particles had been eliminated, CMG particles were gradually removed from the CMGA dataset until a clean set of 15,393 particles produced a map for CMGA with continuous density for the trimeric And1 SepB domain. This set was finally subjected to non-uniform refinement to deliver a final map with a resolution of 6.77 Å.

### Model Building and Refinement

The crystal structures of human GINS (Kamada, Kubota, Arata, Shindo, & Hanaoka, 2007) (PDB DI: 2E9X) and Cdc45 (Simon, Sannino, Costanzo, & Pellegrini, 2016) (5DGO) were docked into the 3.29 Å CMG-DNA map using UCSF Chimera (Pettersen et al., 2004), and manually edited in Coot (Emsley & Cowtan, 2004). Homology models for full-length human MCM2-7 were generated using PHYRE2 (Kelley & Sternberg, 2009), based on the cryo-EM structure of the yeast MCM2-7 double hexamer (3JA8) (N. Li et al., 2015) and docked into the map. Both N-terminal and C-terminal domains of the MCM homology models required extensive rebuilding, which was performed with a combination of remodelling using the Namdinator server (Kidmose et al., 2019) and manual rebuilding. Single-stranded DNA was built in Coot (Emsley & Cowtan, 2004). Models for the ATPase domains of MCM2 and 5, and for the C-terminal domains of MCM2 and 6, were completed using the focussed C-terminal map. The complete CMG-DNA model was subsequently refined using phenix.real_space_refine (Adams et al., 2010) with bond-length and angle restraints for bound ATPγS, magnesium and zinc ions.

For the CMGA structure, the CMG structure and the crystal structure of the AND-1 trimer (5OGS) (Simon et al., 2016) were docked in the map and subjected to rigid-body refinement in phenix.real_space_refine (Adams et al., 2010). The N-tier ring of MCM proteins was treated as a single rigid body, whereas individual ATPase domains in the C-tier were allowed to move independently.

Statistics for real space refinement are reported in Table S2. Figures were prepared using Chimera UCSF (Pettersen et al., 2004) and ChimeraX (Goddard et al., 2018).

### Data availability

CryoEM maps and fitted atomic models have been deposited in the EMDB and PDB. CMG-ATPγS-DNA: EMD-10619, EMD-10620 (C-tier focused map) and PDB ID 6XTX. CMGA: EMD-10621 and PDB ID 6XTY.

## Author contributions

N.J.R. cloned, expressed and purified human CMG and CMGA, reconstituted and determined the cryoEM structure of CMG-ATPγS-DNA and CMGA; S.W.H. and D.Y.C. supervised cryoEM data collections and calculated the cryoEM map of CMGA; V.A.J. helped with purification of CMG; L.P. conceived and supervised the project and wrote the manuscript, with input from N.J.R.

## Acknowledgements

We would like to thank Mairi Kilkenny for help with model building; Christopher Morton for tissue culture support; Laura Radu, Daniel Bollschweiler and Tom Dendooven for advice with cryoEM data processing. This work was funded by a Wellcome Trust investigator award 104641/Z/14/Z to L.P. and by Wellcome Trust awards 202905/Z/16/Z and 206171/Z/17/Z for the establishment of a cryoEM facility in the University of Cambridge.

## References

Abid Ali, F., & Costa, A. (2016). The MCM Helicase Motor of the Eukaryotic Replisome. Journal of molecular biology, 428(9), 1822–1832. Academic Press.

Abid Ali, F., Renault, L., Gannon, J., Gahlon, H. L., Kotecha, A., Zhou, J. C., Rueda, D., et al. (2016). Cryo-EM structures of the eukaryotic replicative helicase bound to a translocation substrate. Nature Communications, 7, 10708.

Adams, P. D., Afonine, P. V., Bunkoczi, G., Chen, V. B., Davis, I. W., Echols, N., Headd, J. J., et al. (2010). PHENIX: a comprehensive Python-based system for macromolecular structure solution. Acta Crystallogr D Biol Crystallogr, 66(Pt 2), 213–221.

Bell, S. P., & Labib, K. (2016). Chromosome Duplication in Saccharomyces cerevisiae. Genetics, 203(3), 1027–1067. Genetics.

Bochman, M. L., Bell, S. P., & Schwacha, A. (2008). Subunit Organization of Mcm2-7 and the Unequal Role of Active Sites in ATP Hydrolysis and Viability. Mol Cell Biol, 28(19), 5865–5873. American Society for Microbiology Journals.

Burnham, D. R., Kose, H. B., Hoyle, R. B., & Yardimci, H. (2019). The mechanism of DNA unwinding by the eukaryotic replicative helicase. Nature Communications, 10(1), 1–14. Nature Publishing Group.

Costa, A., Ilves, I., Tamberg, N., Petojevic, T., Nogales, E., Botchan, M. R., & Berger, J. M. (2011). The structural basis for MCM2-7 helicase activation by GINS and Cdc45. Nature structural & molecular biology, 18(4), 471–477. Nature Publishing Group, a division of Macmillan Publishers Limited. All Rights Reserved.

Douglas, M. E., Ali, F. A., Costa, A., & Diffley, J. F. X. (2018). The mechanism of eukaryotic CMG helicase activation. Nature, 203, 1027. Nature Publishing Group.

Eickhoff, P., Kose, H. B., Martino, F., Petojevic, T., Abid Ali, F., Locke, J., Tamberg, N., et al. (2019). Molecular Basis for ATP-Hydrolysis-Driven DNA Translocation by the CMG Helicase of the Eukaryotic Replisome. Cell reports, 28(10), 2673–2688.e8.

Emsley, P., & Cowtan, K. (2004). Coot: Model-building tools for molecular graphics. Acta Crystallogr D Biol Crystallogr, 60, 2126–2132.

Enemark, E. J., & Joshua-Tor, L. (2006). Mechanism of DNA translocation in a replicative hexameric helicase. Nature, 442(7100), 270–275. Nature Publishing Group.

Fu, Y. V., Yardimci, H., Long, D. T., Ho, T. V., Guainazzi, A., Bermudez, V. P., Hurwitz, J., et al. (2011). Selective bypass of a lagging strand roadblock by the eukaryotic replicative DNA helicase. Cell, 146(6), 931–941. Cell Press.

Gambus, A., Jones, R. C., Sanchez-Diaz, A., Kanemaki, M., van Deursen, F., Edmondson, R. D., & Labib, K. (2006). GINS maintains association of Cdc45 with MCM in replisome progression complexes at eukaryotic DNA replication forks. Nat Cell Biol, 8(4), 358–366. Nature Publishing Group.

Gao, Y., Cui, Y., Fox, T., Lin, S., Wang, H., de Val, N., Zhou, Z. H., et al. (2019). Structures and operating principles of the replisome. Science, 363(6429), eaav7003.

Georgescu, R., Yuan, Z., Bai, L., de Luna Almeida Santos, R., Sun, J., Zhang, D., Yurieva, O., et al. (2017). Structure of eukaryotic CMG helicase at a replication fork and implications to replisome architecture and origin initiation. Proc Natl Acad Sci U S A, 201620500. National Acad Sciences.

Goddard, T. D., Huang, C. C., Meng, E. C., Pettersen, E. F., Couch, G. S., Morris, J. H., & Ferrin, T. E. (2018). UCSF ChimeraX: Meeting modern challenges in visualization and analysis. Protein Science, 27(1), 14–25. John Wiley & Sons, Ltd.

Goswami, P., Abid Ali, F., Douglas, M. E., Locke, J., Purkiss, A., Janska, A., Eickhoff, P., et al. (2018). Structure of DNA-CMG-Pol epsilon elucidates the roles of the non-catalytic polymerase modules in the eukaryotic replisome. Nature Communications, 9(1), 5061.

Ilves, I., Petojevic, T., Pesavento, J. J., & Botchan, M. R. (2010). Activation of the MCM2-7 Helicase by Association with Cdc45 and GINS Proteins. Molecular cell, 37(2), 247–258.

Ishimi, Y. (1997). A DNA helicase activity is associated with an MCM4, -6, and -7 protein complex. Journal of Biological Chemistry, 272(39), 24508–24513. American Society for Biochemistry and Molecular Biology.

Itsathitphaisarn, O., Wing, R. A., Eliason, W. K., Wang, J., & Steitz, T. A. (2012). The hexameric helicase DnaB adopts a nonplanar conformation during translocation. Cell, 151(2), 267–277.

Iyer, L. M., Leipe, D. D., Koonin, E. V., & Aravind, L. (2004). Evolutionary history and higher order classification of AAA+ ATPases. SI:Electron Tomography, 146(1-2), 11– 31.

Kamada, K., Kubota, Y., Arata, T., Shindo, Y., & Hanaoka, F. (2007). Structure of the human GINS complex and its assembly and functional interface in replication initiation. Nature structural & molecular biology, 14(5), 388–396.

Kelley, L. A., & Sternberg, M. J. E. (2009). Protein structure prediction on the Web: a case study using the Phyre server. Nature protocols, 4(3), 363–371.

Kidmose, R. T., Juhl, J., Nissen, P., Boesen, T., Karlsen, J. L., & Pedersen, B. P. (2019). Namdinator - automatic molecular dynamics flexible fitting of structural models into cryo-EM and crystallography experimental maps. IUCrJ, 6(Pt 4), 526–531. International Union of Crystallography.

Kilkenny, M. L., Simon, A. C., Mainwaring, J., Wirthensohn, D., Holzer, S., & Pellegrini, L. (2017). The human CTF4-orthologue AND-1 interacts with DNA polymerase α/primase via its unique C-terminal HMG box. Open Biology, 7(11), 170217. The Royal Society.

Krissinel, E., & Henrick, K. (2007). Inference of macromolecular assemblies from crystalline state. Journal of molecular biology, 372(3), 774–797.

Lam, S. K. W., Ma, X., Sing, T. L., Shilton, B. H., Brandl, C. J., & Davey, M. J. (2013). The PS1 hairpin of Mcm3 is essential for viability and for DNA unwinding in vitro. (R. J. Wellinger, Ed.)PloS one, 8(12), e82177. Public Library of Science.

Langston, L., & O’Donnell, M. (2017). Action of CMG with strand-specific DNA blocks supports an internal unwinding mode for the eukaryotic replicative helicase. eLife, 6, 10708.

Li, N., Zhai, Y., Zhang, Y., Li, W., Yang, M., Lei, J., Tye, B.-K., et al. (2015). Structure of the eukaryotic MCM complex at 3.8 Å. Nature, 524(7564), 186–191.

Liu, X., Schuck, S., & Stenlund, A. (2007). Adjacent residues in the E1 initiator beta-hairpin define different roles of the beta-hairpin in Ori melting, helicase loading, and helicase activity. Molecular cell, 25(6), 825–837.

Lyubimov, A. Y., Strycharska, M., & Berger, J. M. (2011). The nuts and bolts of ring-translocase structure and mechanism. Current Opinion in Structural Biology, 21(2), 240–248. Elsevier Current Trends.

Meagher, M., Epling, L. B., & Enemark, E. J. (2019). DNA translocation mechanism of the MCM complex and implications for replication initiation. Nature Communications, 10(1), 3117. Nature Publishing Group.

Moyer, S. E., Lewis, P. W., & Botchan, M. R. (2006). Isolation of the Cdc45/Mcm2-7/GINS (CMG) complex, a candidate for the eukaryotic DNA replication fork helicase. Proceedings of the National Academy of Sciences of the United States of America, 103(27), 10236–10241.

O’Donnell, Michael E, & Li, H. (2018). The ring-shaped hexameric helicases that function at DNA replication forks. Nature structural & molecular biology, 25(2), 122–130. Nature Publishing Group.

O’Donnell, Michael, Langston, L., & Stillman, B. (2013). Principles and concepts of DNA replication in bacteria, archaea, and eukarya. Cold Spring Harbor perspectives in biology, 5(7), a010108–a010108. Cold Spring Harbor Lab.

Petojevic, T., Pesavento, J. J., Costa, A., Liang, J., Wang, Z., Berger, J. M., & Botchan, M. R. (2015). Cdc45 (cell division cycle protein 45) guards the gate of the Eukaryote Replisome helicase stabilizing leading strand engagement. Proc Natl Acad Sci U S A, 112(3), E249–58.

Pettersen, E. F., Goddard, T. D., Huang, C. C., Couch, G. S., Greenblatt, D. M., Meng, E. C., & Ferrin, T. E. (2004). UCSF Chimera--a visualization system for exploratory research and analysis. J Comput Chem, 25(13), 1605–1612.

Punjani, A., Rubinstein, J. L., Fleet, D. J., & Brubaker, M. A. (2017). cryoSPARC: algorithms for rapid unsupervised cryo-EM structure determination. Nat Methods, 14(3), 290–296. Nature Publishing Group.

Ramey, C. J., & Sclafani, R. A. (2014). Functional conservation of the pre-sensor one beta-finger hairpin (PS1-hp) structures in mini-chromosome maintenance proteins of Saccharomyces cerevisiae and archaea. G3 (Bethesda, Md.), 4(7), 1319–1326. G3: Genes, Genomes, Genetics.

Scheres, S. H. W. (2012). RELION: Implementation of a Bayesian approach to cryo-EM structure determination. SI:Electron Tomography, 180(3), 519–530. Academic Press.

Simon, A. C., Sannino, V., Costanzo, V., & Pellegrini, L. (2016). Structure of human Cdc45 and implications for CMG helicase function. Nature Communications, 7, 11638. Nature Publishing Group.

Simon, A. C., Zhou, J. C., Perera, R. L., van Deursen, F., Evrin, C., Ivanova, M. E., Kilkenny, M. L., et al. (2014). A Ctf4 trimer couples the CMG helicase to DNA polymerase alpha in the eukaryotic replisome. Nature, 510(7504), 293–297.

Singleton, M. R., Sawaya, M. R., Ellenberger, T., & Wigley, D. B. (2000). Crystal structure of T7 gene 4 ring helicase indicates a mechanism for sequential hydrolysis of nucleotides. Cell VL *-*, 101(6), 589–600.

Tegunov, D., & Cramer, P. (2019). Real-time cryo-electron microscopy data preprocessing with Warp. Nat Methods, 16(11), 1146–1152. Nature Publishing Group.

Thomsen, N. D., & Berger, J. M. (2009). Running in reverse: the structural basis for translocation polarity in hexameric helicases. Cell, 139(3), 523–534.

Vijayachandran, L. S., Viola, C., Garzoni, F., Trowitzsch, S., Bieniossek, C., Chaillet, M., Schaffitzel, C., et al. (2011). Robots, pipelines, polyproteins: enabling multiprotein expression in prokaryotic and eukaryotic cells. J Struct Biol, 175(2), 198–208.

Villa, F., Simon, A. C., Ortiz Bazan, M. A., Kilkenny, M. L., Wirthensohn, D., Wightman, M., Matak-Vinkovic, D., et al. (2016). Ctf4 Is a Hub in the Eukaryotic Replisome that Links Multiple CIP-Box Proteins to the CMG Helicase. Molecular cell, 63(3), 385–396.

Yuan, Z., Bai, L., Sun, J., Georgescu, R., Liu, J., O’Donnell, M. E., & Li, H. (2016). Structure of the eukaryotic replicative CMG helicase suggests a pumpjack motion for translocation. Nature structural & molecular biology, 23(3), 217–224. Nature Publishing Group, a division of Macmillan Publishers Limited. All Rights Reserved.

Yuan, Z., Georgescu, R., de Luna Almeida Santos, R., Zhang, D., Bai, L., Yao, N. Y., Zhao, G., et al. (2019). Ctf4 organizes sister replisomes and Pol α into a replication factory. eLife, 8. eLife Sciences Publications Limited.

Zhang, K. (2016). Gctf: Real-time CTF determination and correction. J Struct Biol, 193(1), 1–12.

Zheng, S. Q., Palovcak, E., Armache, J.-P., Verba, K. A., Cheng, Y., & Agard, D. A. (2017). MotionCor2: anisotropic correction of beam-induced motion for improved cryo-electron microscopy. Nat Meth, 14(4), 331–332.

Zivanov, J., Nakane, T., Forsberg, B. O., Kimanius, D., Hagen, W. J., Lindahl, E., & Scheres, S. H. (2018). New tools for automated high-resolution cryo-EM structure determination in RELION-3. eLife, 7.

