## Supplementary information for "CryoEM structures of human CMG - ATPγS - DNA and CMG - AND-1 complexes"

**Table S1.** Data collection parameters for CMG-ATPγS-DNA and CMGA.

| Parameter | CMG-ATPγS -DNA | CMGA |
|---|---|---|
| Detector | Gatan K2 Summit<br>direct electron camera<br>(counting mode) | FEI Falcon3<br>direct electron detector<br>(integrating mode) |
| Nominal magnification | 130,000x | 75,000x |
| No. micrographs | 3,694 | 1,844 |
| Pixel size (Å per pixel)<br>Area ($Å^2$ per pixel) | 1.07<br>1.1449 | 1.06<br>1.1236 |
| Illuminated area (μm) | 1.10 | 1.10 |
| Gun Lens / Spot Size | 4 / 6 | 4 / 3 |
| Energy filter slit (eV) | 20 | N/A |
| Exposure (seconds) | 12 | 0.59 |
| Dose (electrons/pix/sec) | 5.7 | 101.7 |
| Dose (electrons/$Å^2$/sec) | 4.8 | 90.5 |
| No. Detector Frames<br>No. Fractions | 480<br>40 | 23<br>23 |
| Defocus range (μm) | -1.2, -1.5, -1.8, -2.1,<br>-2.4, -2.7 | -1.5, -1.8, -2.1, -2.4, -2.7,<br>-3.0, -3.3 |
| Autofocus | Every 10 μm | Every 10 μm |
| Drift measurement | None | None |
| Delay after stage shift (sec) | 7 | 7 |
| Delay after image shift (sec) | 3 | 3 |
| Exposures per hole | 2 | 2 |
| Objective Aperture | 100 | 100 |
| C2 Aperture | 50 | 70 |

**Table S2.** Real-space refinement.

| | **CMG-ATPγS -DNA** | |
|---|---|---|
| *Model* | | |
| Composition (#) | | |
| Chains | 13 | |
| Atoms | 40829 (Hydrogens: 0) | |
| Residues | Protein: 5058 Nucleotide: 11 | |
| Water | 0 | |
| Ligands | MG: 3 | |
| | ZN: 5 | |
| | AGS: 3 | |
| | ADP: 2 | |
| Bonds (RMSD) | | |
| Length (Å) (#>σ) | 0.009 (1) | |
| Angles (°) (#>σ) | 0.755 (18) | |
| MolProbity score | 2.11 | |
| Clash score | 10.30 | |
| Ramachandran plot (%) | | |
| Outliers | 0.04 | |
| Allowed | 10.89 | |
| Favored | 89.07 | |
| Rotamer outliers (%) | 0.11 | |
| Cβ outliers (%) | 0.00 | |
| Peptide plane (%) | | |
| Cis proline/general | 1.9/0.0 | |
| Twisted proline/general | 0.0/0.0 | |
| ADP (B-factors) | | |
| Iso/Aniso (#) | 40829/0 | |
| min/max/mean | | |
| Protein | 30.19/199.84/91.31 | |
| Nucleotide | 54.57/160.82/85.54 | |
| Ligand | 47.49/210.89/81.95 | |
| Water | --- | |
| Occupancy | | |
| Mean | 1.00 | |
| occ = 1 (%) | 100.00 | |
| 0 < occ < 1 (%) | 0.00 | |
| occ > 1 (%) | 0.00 | |
| *Data* | | |
| Box | | |
| Lengths (Å) | 155.15, 159.43, 208.65 | |
| Angles (°) | 90.00, 90.00, 90.00 | |
| Supplied Resolution (Å) | 3.3 | |
| Resolution Estimates (Å) | Masked | Unmasked |
| d FSC (half maps; 0.143) | --- | --- |
| d 99 (full/half1/half2) | 3.6/---/--- | 3.4/---/--- |
| d model | 3.4 | 3.3 |
| d FSC model (0/0.143/0.5) | 3.1/3.2/3.4 | 3.2/3.3/3.7 |
| Map min/max/mean | -0.04/0.11/-0.00 | |

**Model vs. Data**

====================================================================================

| | |
|---|---|
| CC (mask) | 0.85 |
| CC (box) | 0.81 |
| CC (peaks) | 0.76 |
| CC (volume) | 0.83 |
| Mean CC for ligands | 0.80 |

#### CMGA

*Model*

| | |
|---|---|
| Composition (#) | |
| Chains | 19 |
| Atoms | 48840 (Hydrogens: 0) |
| Residues | Protein: 6117 Nucleotide: 0 |
| Water | 0 |
| Ligands | ZN: 5 |
| Bonds (RMSD) | |
| Length (Å) (#>$\sigma$) | 0.014 (10) |
| Angles (°) (#>$\sigma$) | 1.051 (27) |
| MolProbity score | 2.16 |
| Clash score | 13.26 |
| Ramachandran plot (%) | |
| Outliers | 0.13 |
| Allowed | 9.13 |
| Favored | 90.74 |
| Rotamer outliers (%) | 0.28 |
| Cb outliers (%) | 0.00 |
| Peptide plane (%) | |
| Cis proline/general | 3.8/0.0 |
| Twisted proline/general | 0.0/0.0 |
| ADP (B-factors) | |
| Iso/Aniso (#) | 48840/0 |
| min/max/mean | |
| Protein | 26.78/194.07/81.83 |
| Nucleotide | --- |
| Ligand | 182.70/210.89/198.62 |
| Water | --- |
| Occupancy | |
| Mean | 1.00 |
| occ = 1 (%) | 99.60 |
| 0 < occ < 1 (%) | 0.39 |
| occ > 1 (%) | 0.00 |

*Data*

| | | |
|---|---|---|
| Box | | |
| Lengths (Å) | 160.06, 147.34, 251.22 | |
| Angles (°) | 90.00, 90.00, 90.00 | |
| Supplied Resolution (Å) | 7.0 | |
| Resolution Estimates (Å) | Masked | Unmasked |
| d FSC (half maps; 0.143) | --- | --- |
| d 99 (full/half1/half2) | 8.6/---/--- | 8.3/---/--- |
| d model | 7.2 | 7.2 |
| d FSC model (0/0.143/0.5) | ---/6.6/8.8 | 6.5/6.8/9.2 |
| Map min/max/mean | -0.54/1.64/0.00 | |

*Model vs. Data*

| | |
|---|---|
| CC (mask) | 0.64 |
| CC (box) | 0.81 |
| CC (peaks) | 0.58 |
| CC (volume) | 0.63 |
| Mean CC for ligands | 0.40 |

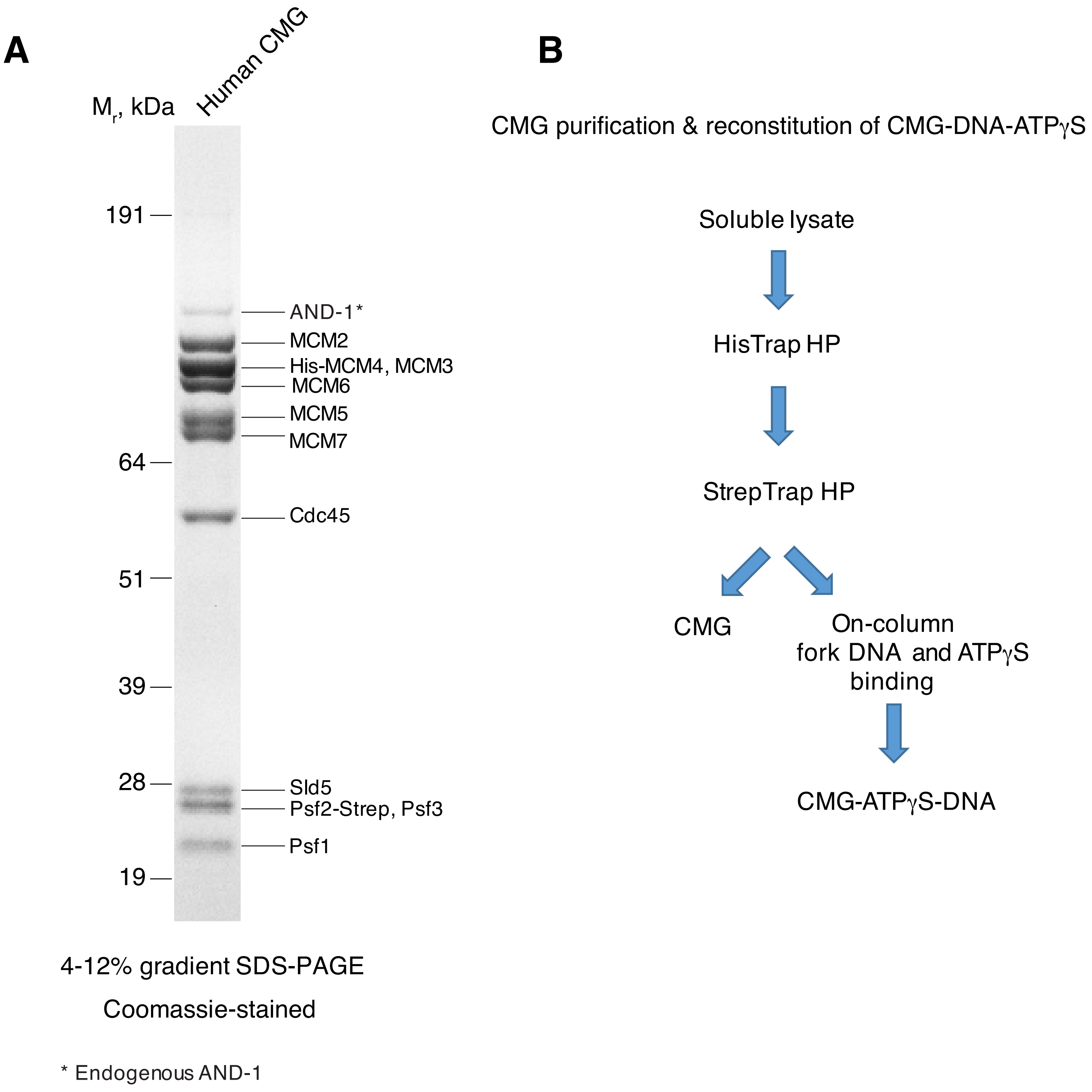


**Supplementary figure 1**

### Supplementary figure 2

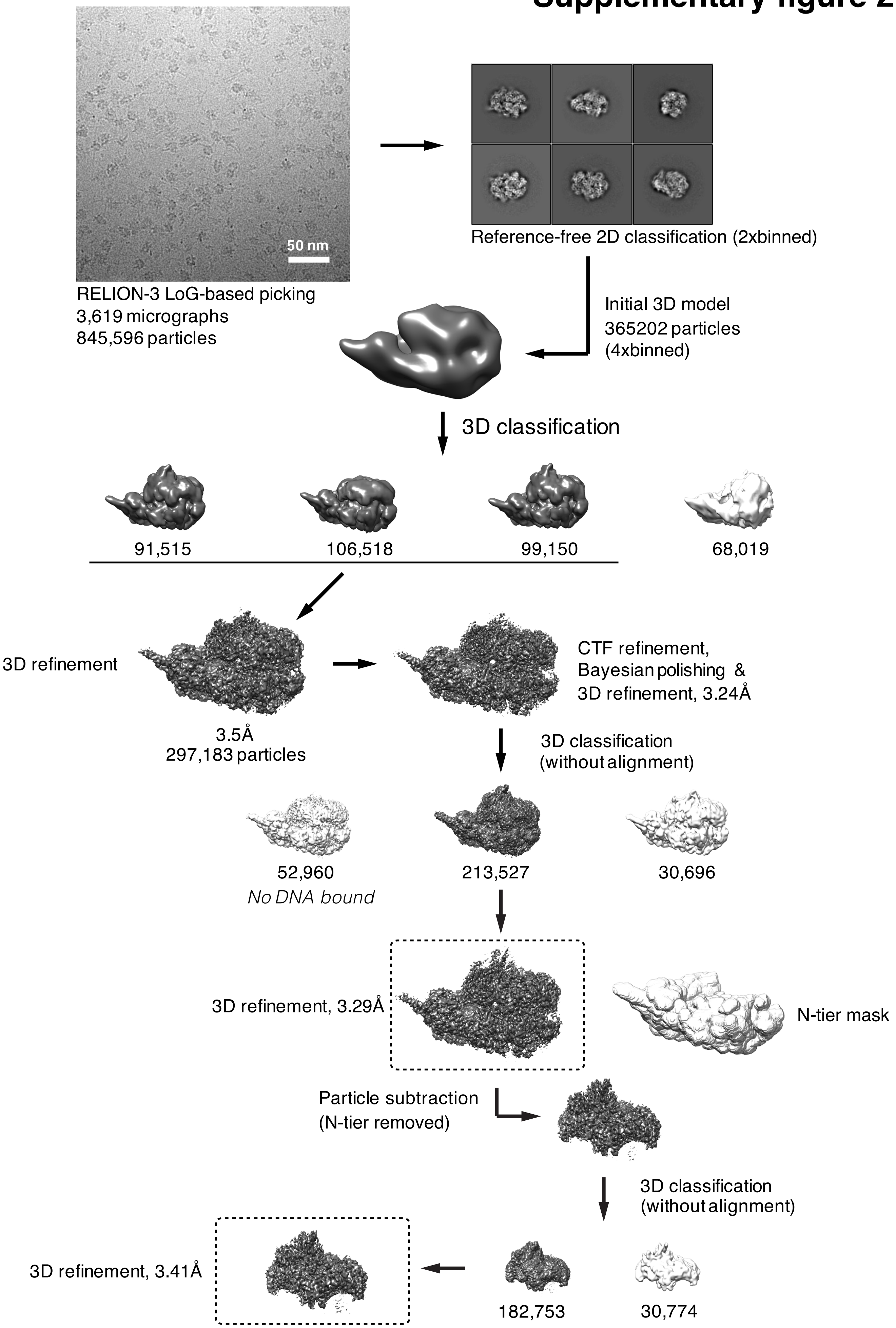

**A**

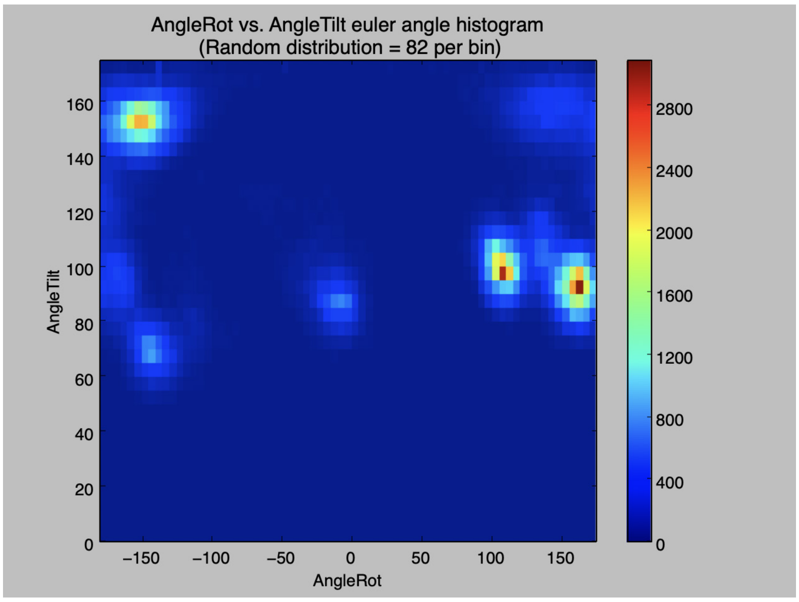


**B**

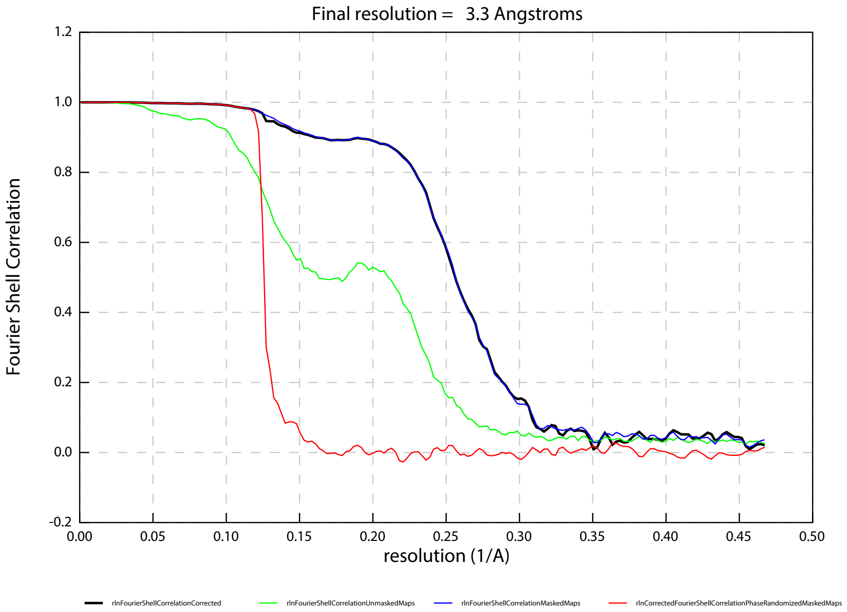


### Supplementary figure 3

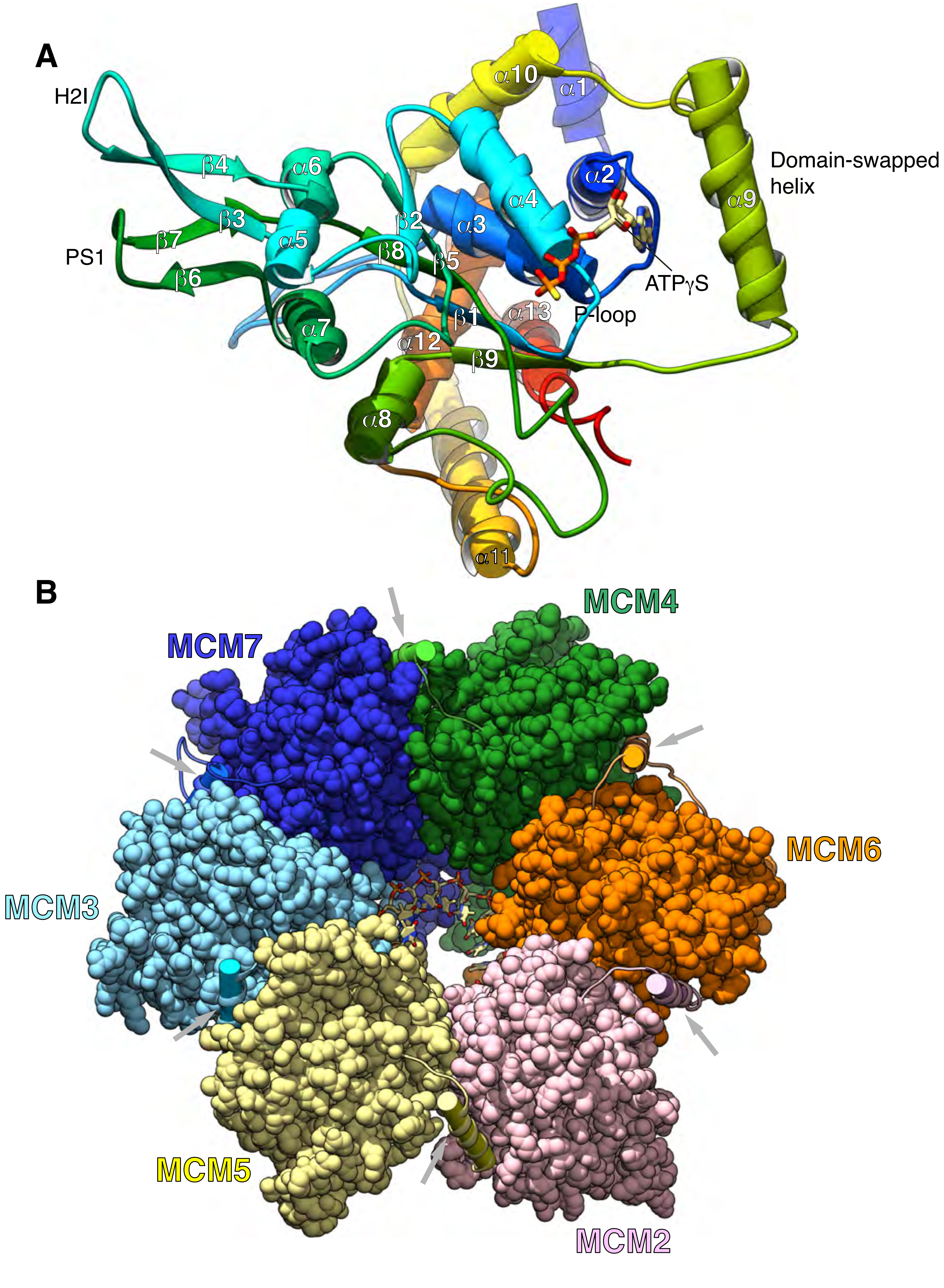


**Supplementary figure 4**

A

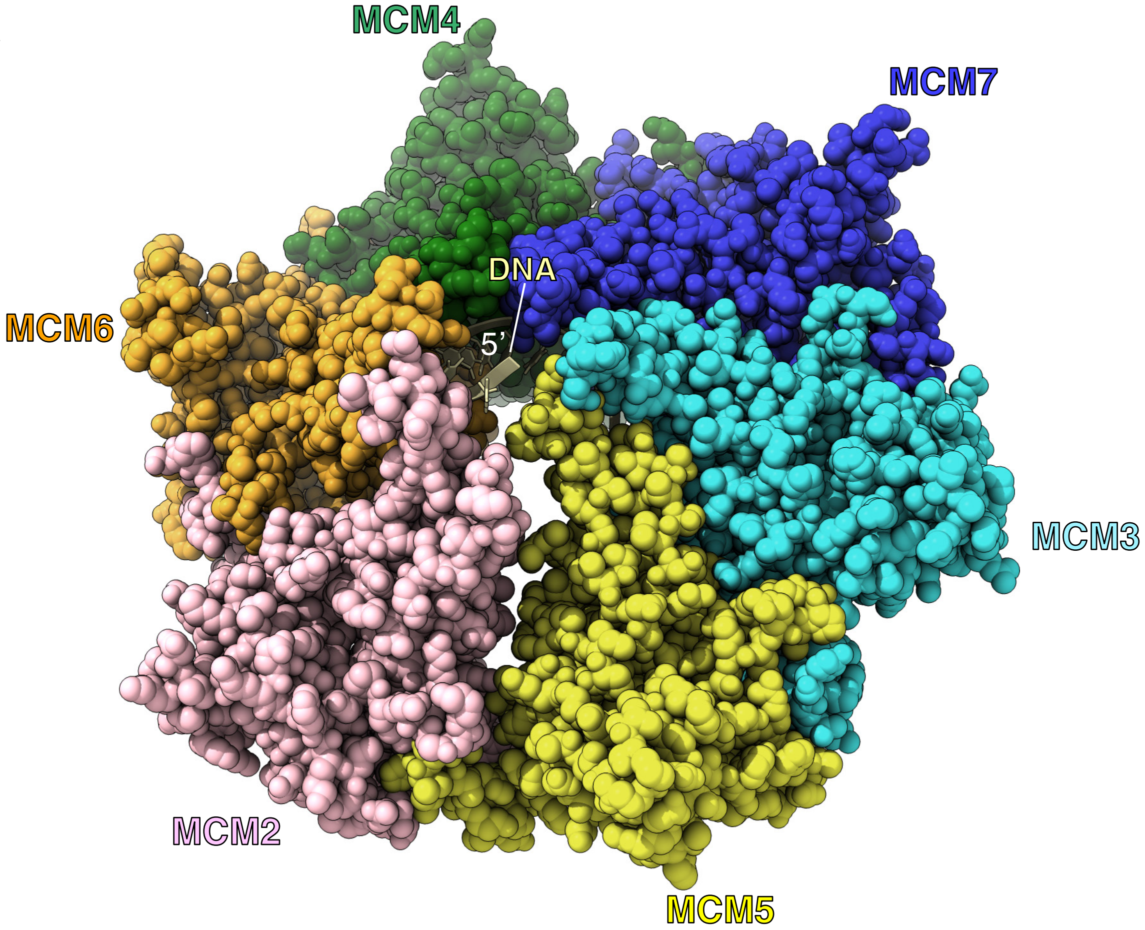


B

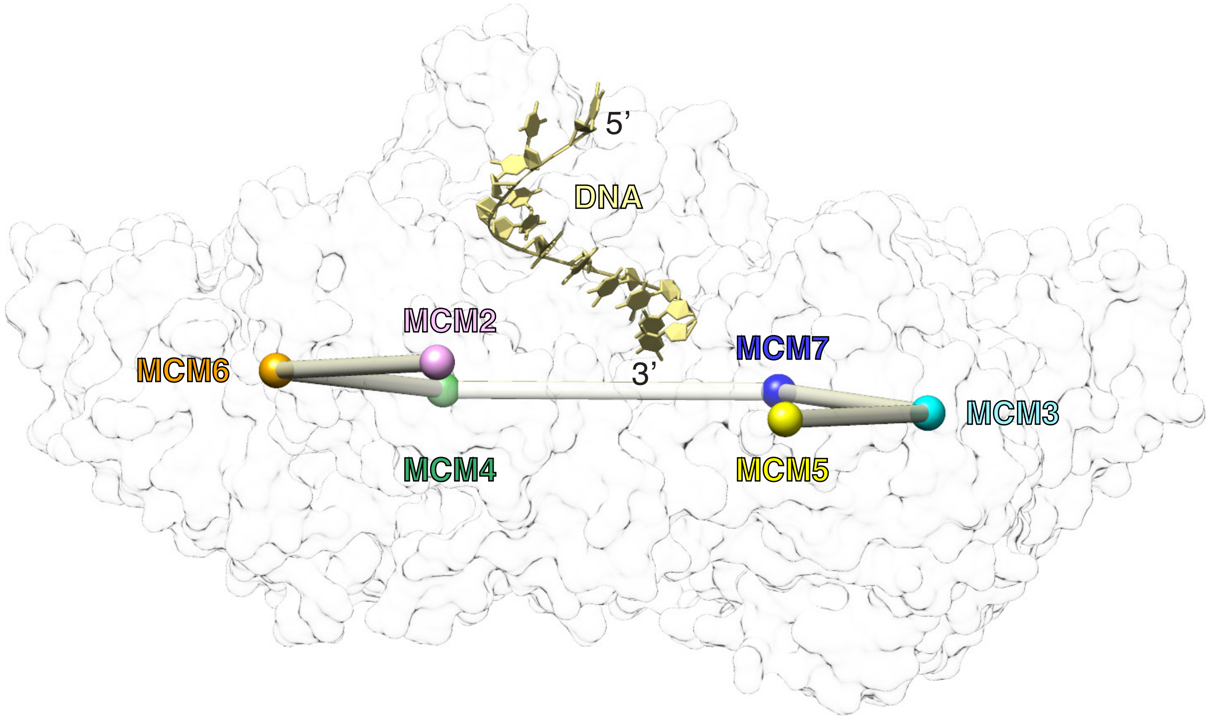


**Supplementary figure 5**

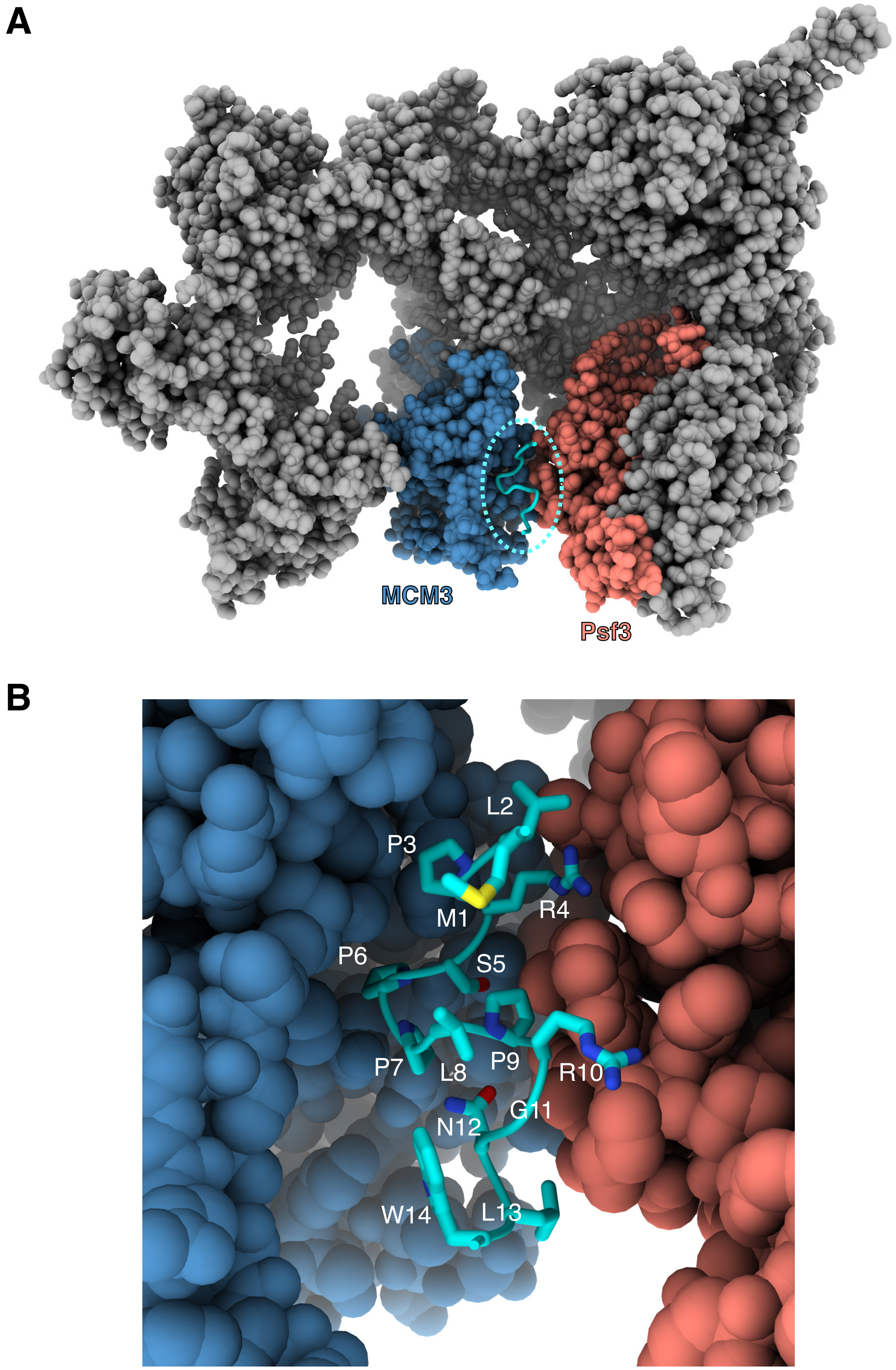


**Supplementary figure 6**

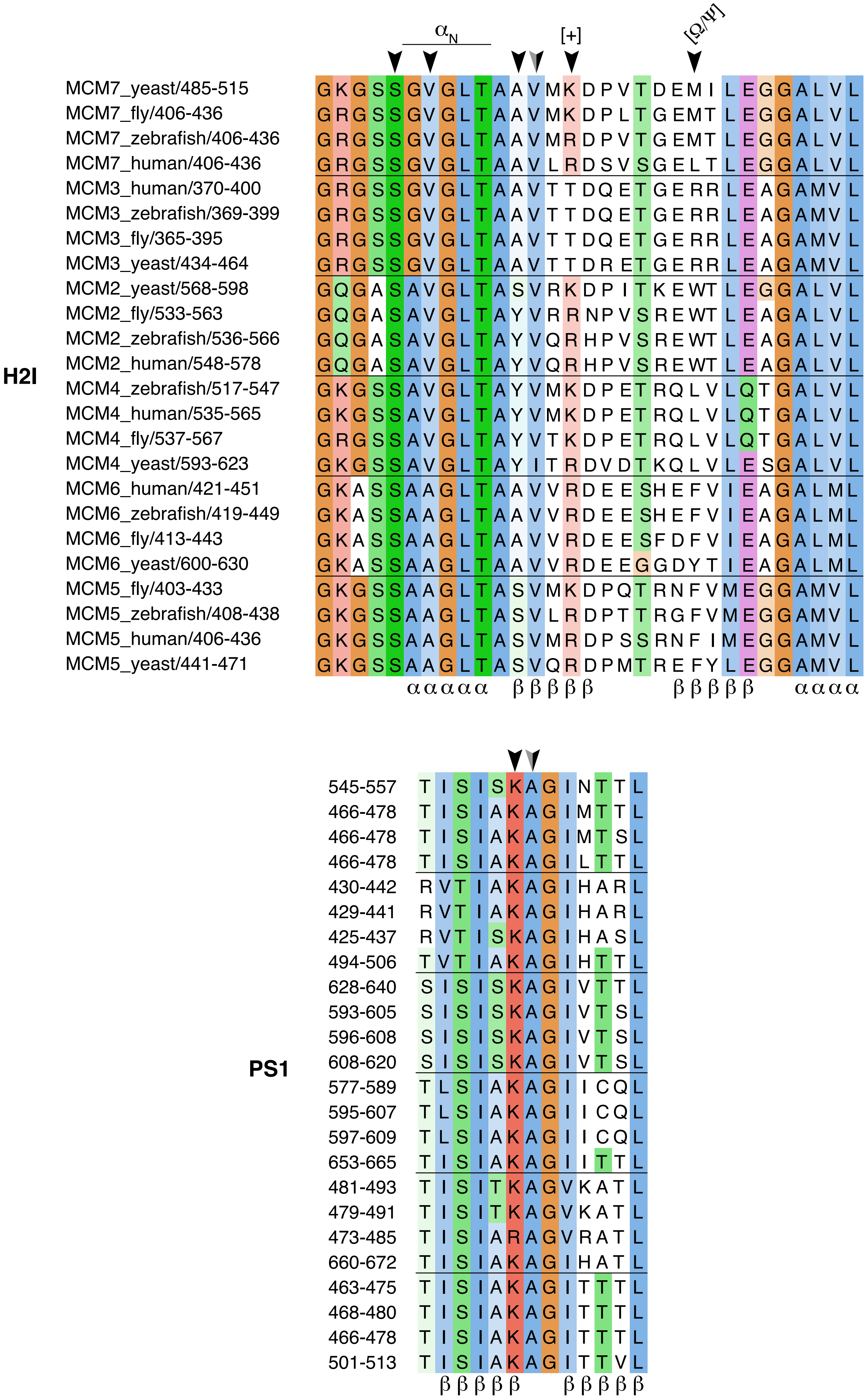


**Supplementary figure 7**

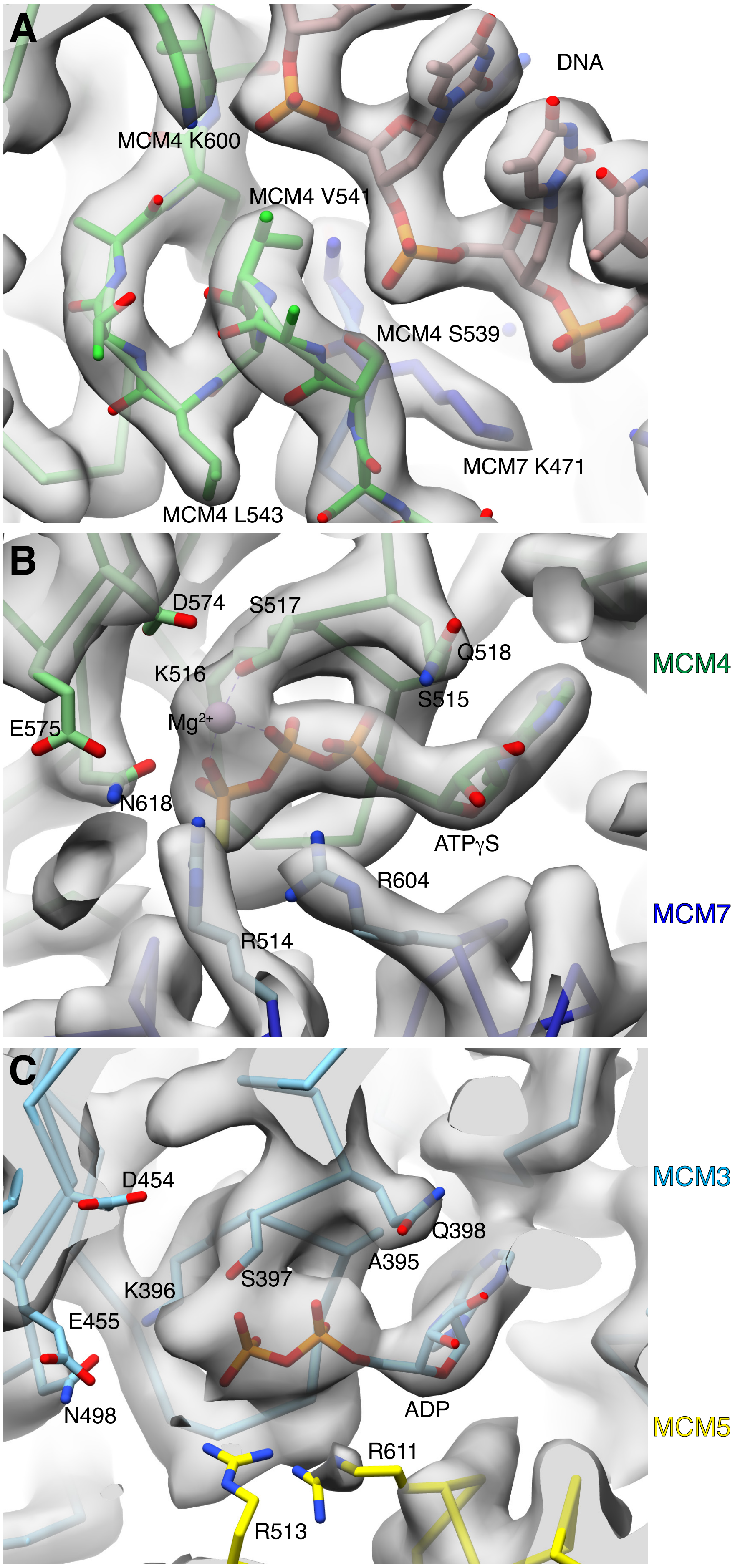


**Supplementary figure 8**

| | Walker A | | Walker B |
|---|---|---|---|
| MCM7_yeast/460-468 | GDPGVAKSQ | 519-525 | GICCIDE |
| MCM7_fly/381-389 | GDPGVAKSQ | 440-446 | GVCCIDE |
| MCM7_zebrafish/381-389 | GDPGVAKSQ | 440-446 | GVCCIDE |
| MCM7_human/381-389 | GDPGVAKSQ | 440-446 | GVCCIDE |
| MCM3_human/345-353 | GDPSVAKSQ | 404-410 | GVVCIDE |
| MCM3_zebrafish/344-352 | GDPSVAKSQ | 403-409 | GVVCIDE |
| MCM3_fly/340-348 | GDPSVAKSQ | 399-405 | GVVCIDE |
| MCM3_yeast/409-417 | GDPSTAKSQ | 468-474 | GVVCIDE |
| MCM2_yeast/543-551 | GDPGTAKSQ | 602-608 | GVCLIDE |
| MCM2_fly/508-516 | GDPGTAKSQ | 567-573 | GVCLIDE |
| MCM2_zebrafish/511-519 | GDPGTAKSQ | 570-576 | GVCLIDE |
| MCM2_human/523-531 | GDPGTAKSQ | 582-588 | GVCLIDE |
| MCM4_zebrafish/492-500 | GDPGTSKSQ | 551-557 | GICCIDE |
| MCM4_human/510-518 | GDPGTSKSQ | 569-575 | GICCIDE |
| MCM4_fly/512-520 | GDPGTSKSQ | 571-577 | GVCCIDE |
| MCM4_yeast/568-576 | GDPSTSKSQ | 627-633 | GVCCIDE |
| MCM6_human/396-404 | GDPSTAKSQ | 455-461 | GVCCIDE |
| MCM6_zebrafish/394-402 | GDPSTAKSQ | 453-459 | GVCCIDE |
| MCM6_fly/388-396 | GDPSTAKSQ | 447-453 | GICCIDE |
| MCM6_yeast/575-583 | GDPSTSKSQ | 634-640 | GICCIDE |
| MCM5_fly/378-386 | GDPGTAKSQ | 437-443 | GVVCIDE |
| MCM5_zebrafish/383-391 | GDPGTAKSQ | 442-448 | GVVCIDE |
| MCM5_human/381-389 | GDPGTAKSQ | 440-446 | GVVCIDE |
| MCM5_yeast/416-424 | GDPGTAKSQ | 475-481 | GVVCIDE |

| | Sensor-1 * | | Sensor-3 * | | Arg finger * | | Sensor-2 * |
|---|---|---|---|---|---|---|---|
| 567-569 | ANP | 537-543 | IHEVMEQ | 591-594 | LSRF | 685-690 | TPRTLL |
| 488-490 | ANP | 458-464 | IHEVMEQ | 512-515 | LSRF | 602-607 | SARNLL |
| 488-490 | ANP | 458-464 | IHEVMEQ | 512-515 | LSRF | 602-607 | SARTLL |
| 488-490 | ANP | 458-464 | IHEVMEQ | 512-515 | LSRF | 602-607 | SARTLL |
| 452-454 | ANP | 422-428 | IHEVMEQ | 476-479 | LSRF | 614-619 | TARTLE |
| 451-453 | ANP | 421-427 | IHEVMEQ | 475-478 | LSRF | 613-618 | TARTLE |
| 447-449 | ANP | 417-423 | IHEVMEQ | 471-474 | LSRF | 608-613 | TARTLE |
| 516-518 | ANP | 486-492 | IHEVMEQ | 540-543 | LSRF | 698-703 | TARTLE |
| 650-652 | ANP | 620-626 | IHEAMEQ | 674-677 | LSRF | 806-811 | TVRHLE |
| 615-617 | ANP | 585-591 | IHEAMEQ | 639-642 | LSRF | 740-745 | TVRHIE |
| 618-620 | ANP | 588-594 | IHEAMEQ | 642-645 | ISRF | 748-753 | TVRHIE |
| 630-632 | ANP | 600-606 | IHEAMEQ | 654-657 | ISRF | 763-768 | TVRHIE |
| 599-601 | ANP | 569-575 | LHEVMEQ | 623-626 | LSRF | 712-717 | YPRQLE |
| 617-619 | ANP | 587-593 | LHEVMEQ | 641-644 | LSRF | 730-735 | YPRQLE |
| 619-621 | ANP | 589-595 | LHEVMEQ | 643-646 | LSRF | 732-737 | YPRQLE |
| 675-677 | ANP | 645-651 | LHEVMEQ | 699-702 | LSRF | 794-799 | TTRQLE |
| 503-505 | ANP | 473-479 | IHEAMEQ | 527-530 | MSRF | 617-622 | TVRQLE |
| 501-503 | ANP | 471-477 | IHEAMEQ | 525-528 | MSRF | 615-620 | TVRQLE |
| 495-497 | ANP | 465-471 | IHEAMEQ | 519-522 | MSRF | 609-614 | TVRQLE |
| 682-684 | ANP | 652-658 | IHEAMEQ | 706-709 | MSRF | 796-801 | TVRQLE |
| 485-487 | ANS | 455-461 | IHEAMEQ | 508-511 | LSRF | 608-613 | TVRQLE |
| 490-492 | ANS | 460-466 | IHEAMEQ | 513-516 | LSRF | 611-616 | TVRQLE |
| 488-490 | ANS | 458-464 | IHEAMEQ | 511-514 | LSRF | 609-614 | TVRQLE |
| 523-525 | ANP | 493-499 | IHEAMEQ | 547-550 | LSRF | 649-654 | TIRQLE |

**Supplementary figure 9**

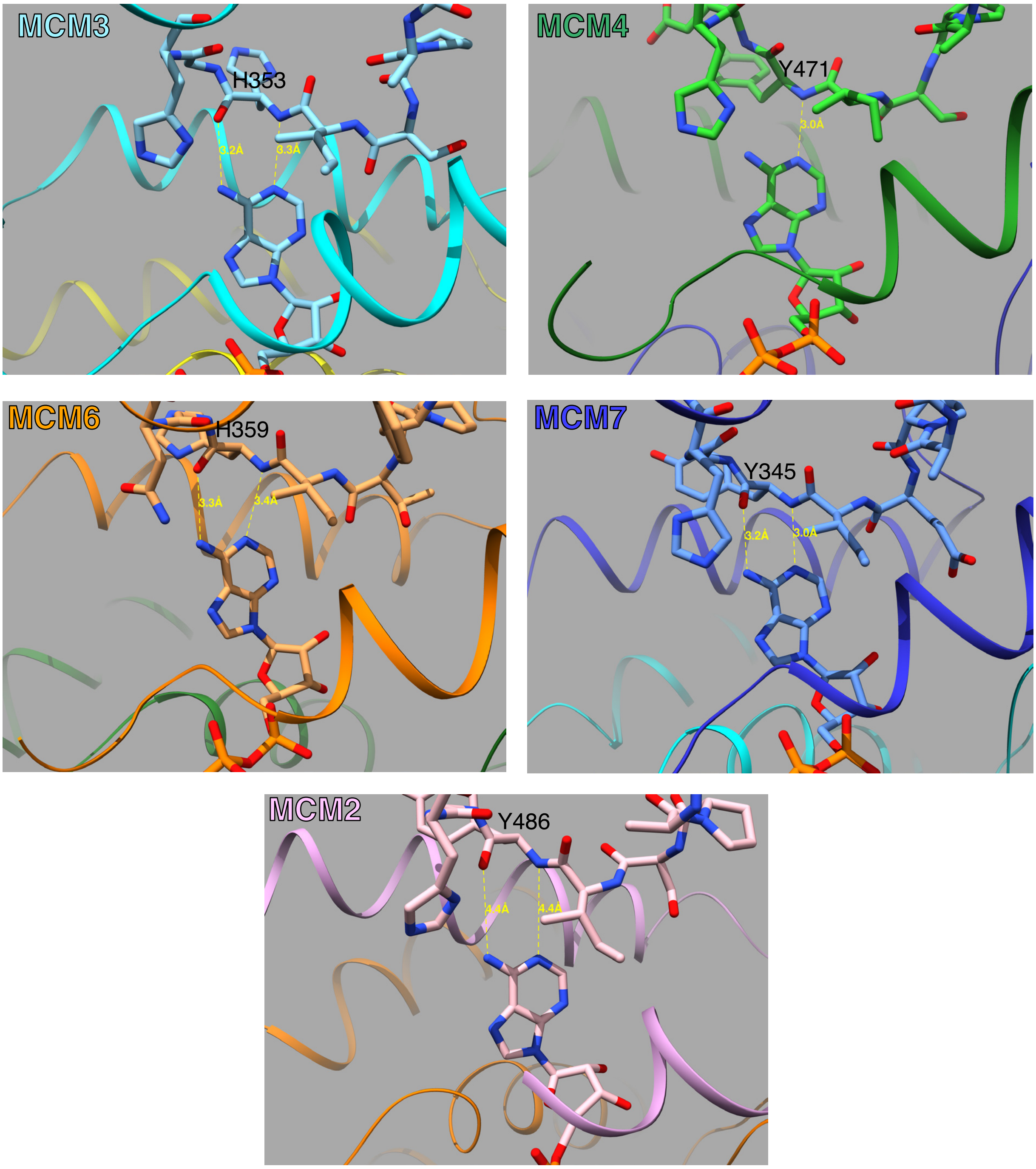


**Supplementary figure 10**

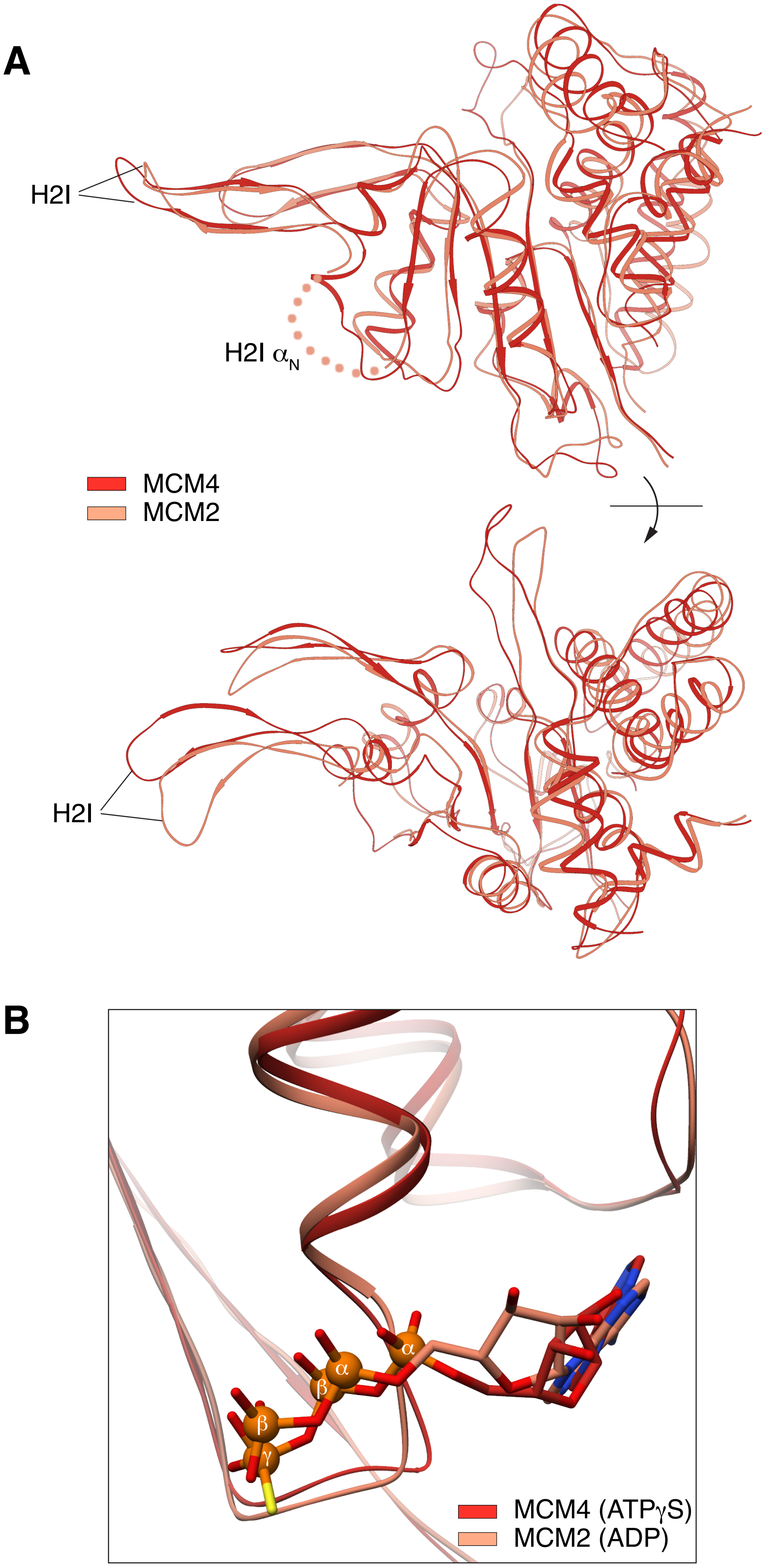


**Supplementary figure 11**

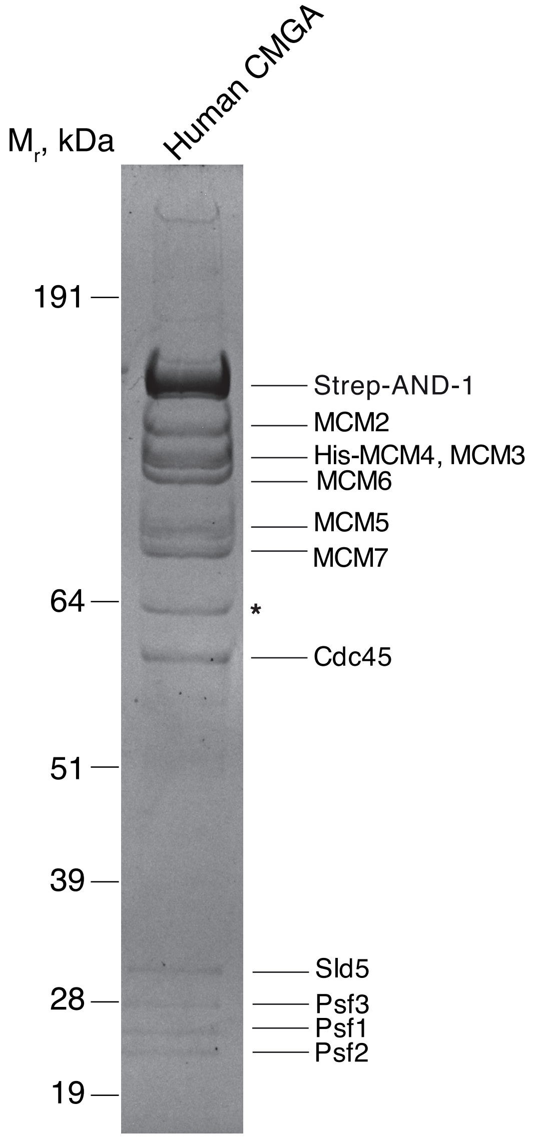


4-12% gradient SDS-PAGE

Coomassie-stained

**Supplementary figure 12**

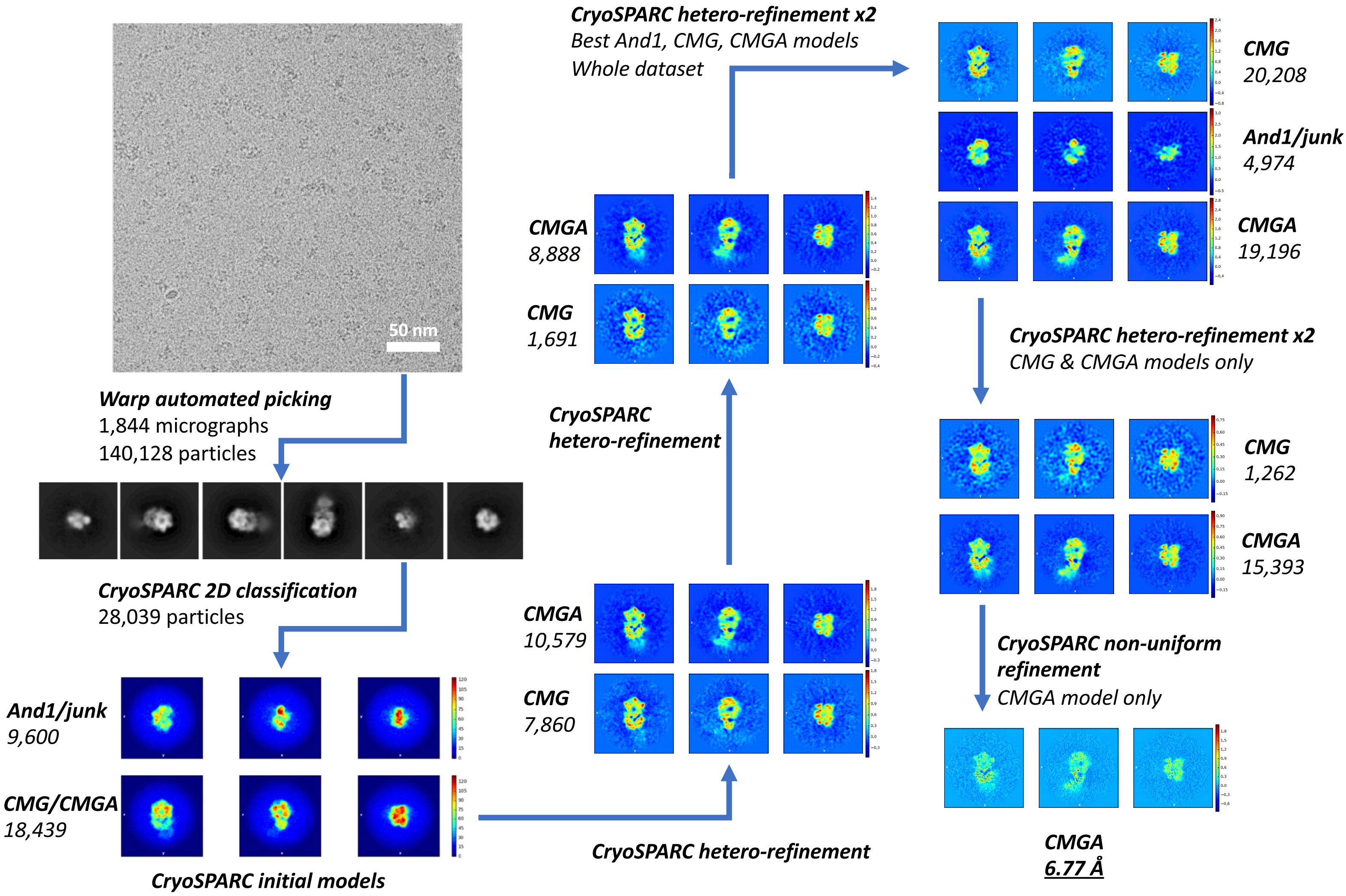


### Supplementary figure 13

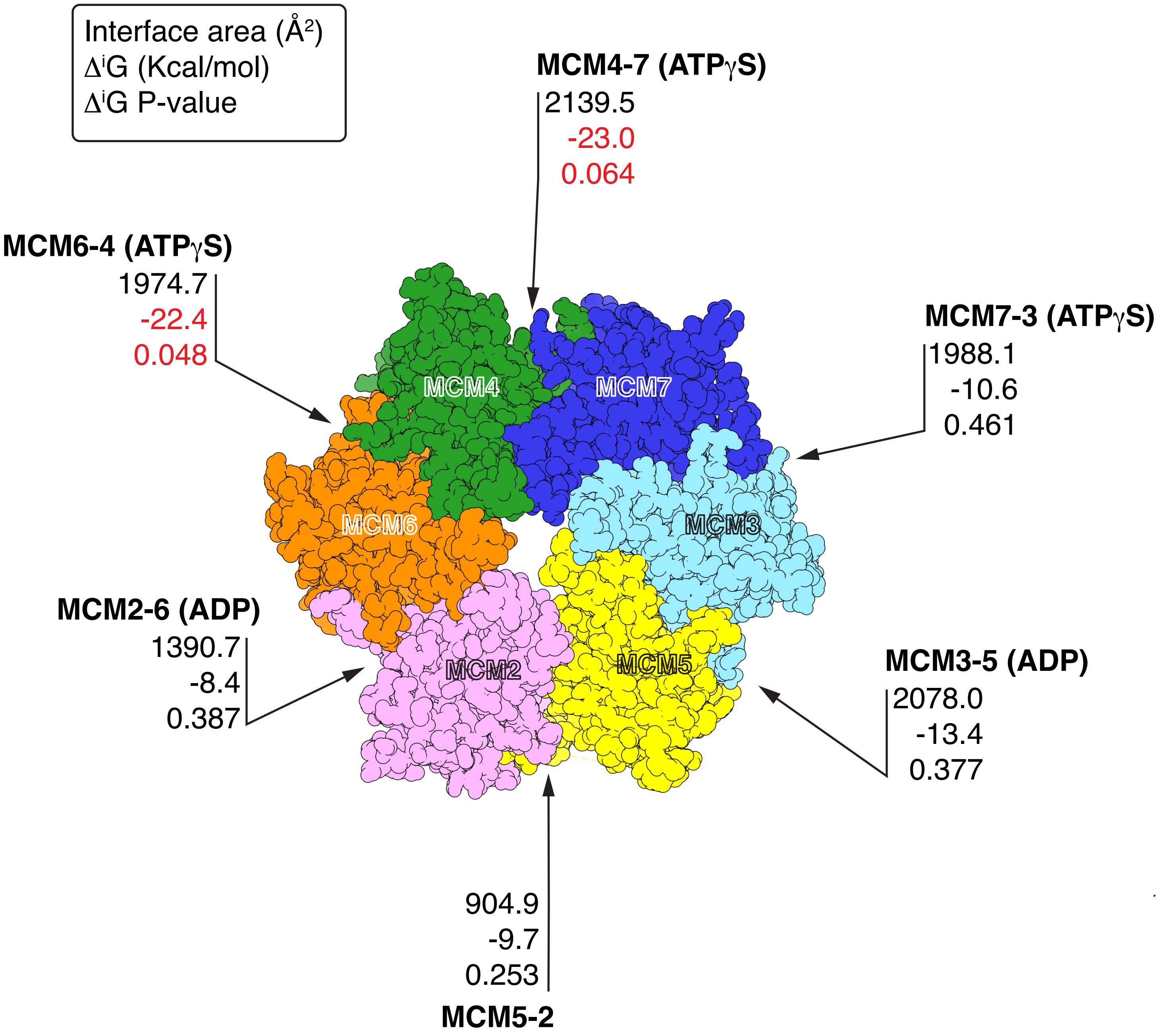


**Supplementary figure 14**

**Supplementary Figure Legend**

**Figure S1.** Purification of recombinant human CMG produced by transient transfection in HEK293 Free Style™ cells. **A** SDS-PAGE analysis of purified CMG. **B** Schematic protocol for purification of CMG and CMG-ATPγS-DNA.

**Figure S2.** CryoEM data processing and structure determination in Relion (Scheres, 2012) for CMG-ATPγS-DNA. 3D classes in white were not considered for further processing.

**Figure S3.** Statistical analysis of cryoEM data processing. **A** Heat map of CMG-ATPγS-DNA particle distribution. The diagram was generated using scripts from the Leschziner lab (github.com/leschzinerlab/Relion). **B** Fourier Shell Correlation curve, generated using PostProcessing in Relion (Scheres, 2012).

**Figure S4.** Secondary structure elements of the AAA+ ATPase domain of human MCM proteins. **A** Cartoon drawing of MCM7 ATPase domain, chosen as representative of the six MCM proteins. The protein is coloured N-end (blue) to C-end (red). α-helices are shown as cylinders (α1- α10), and β-strands (β1- β9) as arrows. Secondary structure elements and notable structural element of the ATPase domain are annotated. **B** Spacefill representation of the MCM C-tier, coloured as in Figure 1. The domain-swapped helix is shown as a cylinder and its position indicated by a grey arrow.

**Figure S5.** Conformation of the MCM C-tier. **A** Spacefill representation of the MCM C-tier. The view highlights the open MCM2-5 interface. The MCM subunits are coloured as in Figure 1. **B** The MCM C-tier ring departs slightly from planarity. The position of each MCM ATPase domain is represented by a sphere located at its centre of mass, calculated in Chimera UCSF (Pettersen et al., 2004). The spheres are coloured according to MCM subunit and neighbouring spheres are joined by a stick. The C-tier is shown as a transparent white surface and the DNA as a ribbon drawing in khaki.

**Figure S6.** The N-tail of MCM3 augments the interface between the MCM N-tier and GINS. **A** Spacefill view of the MCM N-tier - GINS interaction. The N-tail of MCM3 is shown as cyan tube, MCM3 is in light blue and the Psf3 subunit of GINS is in salmon. **B** Close-up view of MCM3-GINS interface. The side chains of the amino acids in the MCM3 N-tail are shown as sticks.

**Figure S7.** Multiple sequence alignment of the H2I (top) and PS1 (bottom) DNA-binding loops of the MCM2-7 ATPase domains. For each MCM protein, the homologous sequences for the yeast, fly, zebrafish and human proteins are shown. The residue conservation in the alignment is coloured according to the Clustal colour scheme. The amino acids that interact with the DNA are indicated by black arrowheads. Mixed grey-black arrowheads indicate residues that use a main-chain nitrogen to hydrogen bond a phosphate group in the DNA. The extent of the H2I $\alpha_N$ helix and the position of the conserved DNA-binding residues [+] and [Ω/Ψ] are indicated. The secondary structure elements are also marked below the alignment.

**Figure S8.** Details of the cryoEM map for human CMG-ATPγS-DNA. The map is shown as a transparent grey surface, the protein and DNA are drawn as sticks. The MCM proteins are coloured as in Figure 1. Relevant amino acids are labelled. **A** The MCM-DNA interface. **B** The ATP-binding site at the MCM4-7 interface, with bound ATPγS. **C** The ATP-binding site at the MCM3-5 interface, with bound ADP.

**Figure S9.** Multiple sequence alignment of the ATP-binding motifs in the MCM2-7 ATPase domains. For each MCM protein, the homologous sequences for the yeast, fly, zebrafish and human proteins are shown. The residue conservation in the alignment is coloured according to the Clustal colour scheme. The position of the sensor amino acid in each of the Sensor 1-3 and 'Arg finger' motifs is marked with an asterisk.

**Figure S10.** Watson and Crick-like hydrogen bonding of the adenine base in ATPγS and ADP with the main-chain nitrogen and carbonyl oxygen of the opposing amino acid. Adoption of a conformation suitable for hydrogen bonding with the carbonyl group requires a conserved glycine following the amino acid bound to the adenine; the glycine in MCM4 is replaced with a glutamate and therefore only one hydrogen bond is observed. The position of ADP in MCM2 does not allow for hydrogen bonding formation, as the adenine is too far from the protein.

**Figure S11.** DNA- and ADP-binding features of the MCM2 subunit. The ATPase domain of MCM2 is superimposed to that of MCM4 for comparison. The proteins are shown as thin ribbons and coloured red (MCM4) and pink (MCM2). **A** Two views at 90° of the MCM2 and MCM4 ATPase domains. In the top view, the H2I $\alpha_N$ helix of MCM2 is disordered and drawn as a dotted line. **B** The ATP-binding sites of MCM2 and MCM4, with bound ADP and ATPγS respectively. The positions of the α and β phosphates of the ADP and of the α, β and γ phosphate of ATPγS are highlighted by small spheres.

**Figure S12.** SDS-PAGE of purified recombinant human CMGA (Cdc45-MCM-GINS-AND-1) produced by transient transfection in HEK293 Free Style™ cells. The asterisk indicates an unidentified protein contaminant.

**Figure S13.** CryoEM data processing and structure determination in CryoSparc (Punjani, Rubinstein, Fleet, & Brubaker, 2017) for CMGA.

**Figure S14.** Solvation free-energy analysis of interface formation in the MCM2-7 C-tier ring. MCM subunits are shown in spacefill representation, and colour coded as in Figure 1. For each interface, the values of buried interface area, $\Delta^{i}G$ (Kcal/mol) and $\Delta^{i}G$ P-value are shown. The analysis was performed using the PISA server at EBI (https://www.ebi.ac.uk/pdbe/pisa/). The $\Delta^{i}G$ values for the MCM6-4 and MCM4-7 interfaces are highlighted in red.


Pettersen, E. F., Goddard, T. D., Huang, C. C., Couch, G. S., Greenblatt, D. M., Meng, E. C., & Ferrin, T. E. (2004). UCSF Chimera--a visualization system for exploratory research and analysis. *J Comput Chem*, *25*(13), 1605–1612.

Punjani, A., Rubinstein, J. L., Fleet, D. J., & Brubaker, M. A. (2017). cryoSPARC: algorithms for rapid unsupervised cryo-EM structure determination. *Nat Methods*, *14*(3), 290–296. Nature Publishing Group.

Scheres, S. H. W. (2012). RELION: Implementation of a Bayesian approach to cryo-EM structure determination. *SI:Electron Tomography*, *180*(3), 519–530. Academic Press.
